# Structural basis for ATP regulation of human 5-lipoxygenase

**DOI:** 10.1101/2025.11.26.690380

**Authors:** Tarvi Teder, Federico Pozzani, Tim Schulte, A. Najla Hosseini, Olof Rådmark, Laura Orellana, Jesper Z. Haeggström

**Affiliations:** Department of Medical Biochemistry and Biophysics, Division of Chemistry II, Karolinska Institutet, Solnavägen 9, SE-171 65 Stockholm, Sweden; Department of Oncology-Pathology, Karolinska Institutet, SE-171 77 Stockholm, Sweden; Department of Biochemistry and Biophysics, National Bioinformatics Infrastructure Sweden, Science for Life Laboratory, Stockholm University, SE-171 21 Solna, Sweden

## Abstract

5-Lipoxygenase catalyzes the committed step in the biosynthesis of the powerful proinflammatory leukotrienes and pro-resolving anti-inflammatory lipoxins. Here we present a 3.3 Å cryo-EM structure of wild type 5-lipoxygenase in complex with ATP, one of its most important allosteric regulators. The nucleotide is located in a positively charged pocket of the catalytic domain and held in place by a complex network of amino acid side chains and backbone carbonyl and amino groups. Mutagenetic analysis suggests that ATP binding and action is primarily mediated via Lys320, assisted by Gln657. Further atomistic simulations demonstrate that ATP binding induces movements of the PLAT domain coupled to conformational rearrangements at the active center. These findings provide important structural and regulatory insights to a key enzyme in leukotriene biosynthesis, thus aiding in design of new antiphlogistic drugs.

## INTRODUCTION

Leukotrienes are potent lipid mediators in inflammatory conditions such as asthma, rhinitis and cardiovascular diseases, while lipoxins and related compounds have anti-inflammatory and pro-resolving properties^1, 2^. 5-Lipoxygenase (5-LOX) is the central enzyme in the leukotriene and lipoxin cascades where it oxidizes arachidonic acid into 5*S*-hydro(pero)xy eicosatetraenoic acid (5*S*-H(p)ETE), which is further dehydrated to yield the unstable epoxide intermediate leukotriene A_4_. Unlike other members of the lipoxygenase family, 5-LOX is subject to complex regulation by phosphorylation, membrane association, and several soluble allosteric factors, most notably calcium and ATP. In intact immune cells, 5-LOX traffics from the cytosol or nucleosol to the perinuclear membrane, where it meets up with its partner protein, FLAP, which is required for maximal activity and leukotriene output.

Due to inherent instability, 5-LOX escaped structural characterization for decades until Newcomer and coworkers succeeded in crystallizing a variant of 5-LOX engineered to improve stability and solubility. The structure of this mutated protein, named Stable 5-LOX, was solved at 2.4 Å resolution^3^.

ATP is one of the classical factors known to stimulate 5-LOX and is required to reach maximal enzyme activity *in vitro*^4^. The EC_50_ of ATP has been estimated to ~30 µM and both the initial hydroperoxidation of arachidonic acid (AA) and the following epoxidation into LTA_4_, are stimulated by the nucleotide^5, 6^. The mechanism of 5-LOX activation by ATP is not known but involves binding of the nucleotide to a presumed allosteric site in the protein without any apparent hydrolysis of phosphodiester bonds. Here, we determined the cryo-EM structure of wildtype 5-LOX at 3.3 Å resolution in complex with ATP. Together with biochemical and computational evidence the structure reveals the molecular basis for ATP regulation of 5-LOX and offers insights to the mechanism for this allosteric enzyme activator.

## RESULTS AND DISCUSSION

### Cryo-EM structure of wild type 5-LOX in complex with ATP

Single-particle cryo-EM data were collected for wild-type 5-LOX prepared in the presence of ATP, yielding a high-quality 3.3 Å resolution map (Table 1, Supplementary Table 1 and Supplementary Fig. 1) with excellent map support for 90% of the model (Fig. 1A and 1B). Overall, the wild-type 5-LOX structure is virtually identical to the crystal structure of Stable 5-LOX^3^, exhibiting a Cα RMSD-value of 0.675 Å. 5-LOX exhibits a typical lipoxygenase structure with the N-terminal residues 1-116 folded into a C2-like, so called PLAT (Polycystin-1, Lipoxygenase, Alpha-Toxin) domain, also referred to as the β-barrel domain, fused to a larger catalytic domain harboring the non-heme iron (Fig. 1C). Five major cavities were found in wild-type 5-LOX (Supplementary Fig. 3), one of which (pocket 1, P1) likely binds AA (Fig. 1E-F).

**Figure 1.**
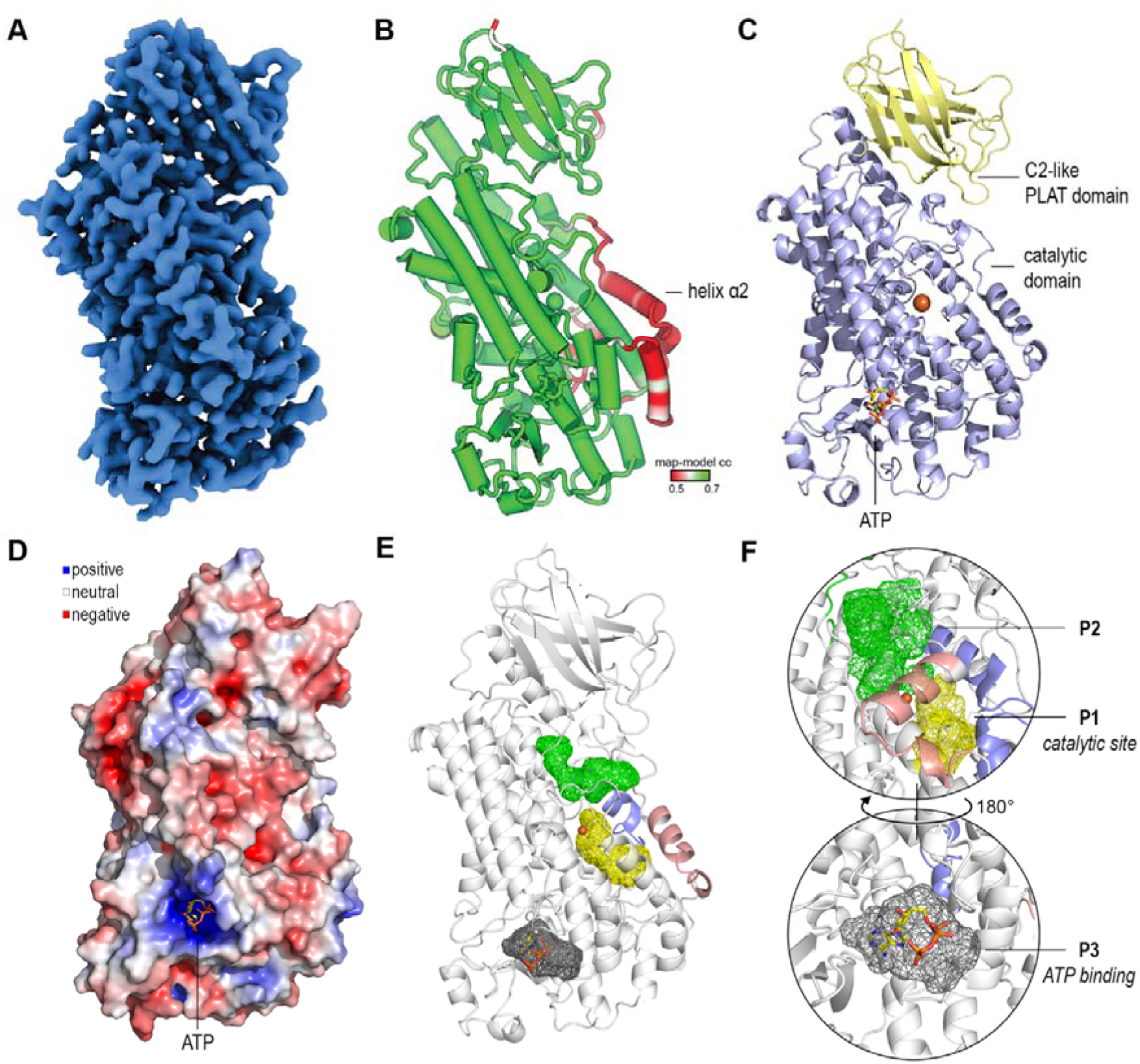
Cryo-EM structure of wild-type 5-LOX in complex with ATP. **A** The density map of 5-LOX is presented in blue. **B** The model of 5-LOX is colored in green, white and red for map-model correlation values of 0.7, 0.6, and 0.5, respectively. Thus, α2H as well as parts of the arched helix lack map support. **C** The model of 5-LOX resolved at 3.3 Å showing the catalytic domain (purple) and C2-like PLAT domain (yellow). The bound ATP molecule is depicted in stick representation with carbon atoms in yellow, phosphorus in orange, nitrogen in blue and oxygen in red. **D** The 5-LOX protein surface is colored using a red (negative), white (neutral) and blue (positive) color gradient for Coulombic electrostatic potential (ESP) in the range of −5 to +5 kcal/(mol·e), respectively. The bound ATP molecule is depicted in stick representation with carbon atoms in yellow, phosphorus in orange, nitrogen in blue and oxygen in red. **E, F** Solvent-accessible pockets in the catalytic domain (P1–P3) were identified in the 5-LOX structure using PyVOL with volume thresholds of 1.5–4.0 Å³. P1 corresponds to the catalytic site (yellow), while P3 represents the ATP-binding pocket (grey). The α2H and arched helix covering the catalytic site (P1) are highlighted in pink and light purple, respectively. The pocket P2 (green) is of unknown function. See related Supplementary Fig. 3, containing additional pockets and docking simulations with the AA substrate.

**Table 1.** Refinement and validation of Cryo-EM data.

|  |  |
| --- | --- |
| Map resolution (FSC 0.143, masked) | 3.3 Å |
| Map sharpening B-factor (Å<sup>2</sup>) | -110 |
| Model-map CC (mask) | 0.84 |
| RMSD bonds (Å) / angles (°) | 0.008 / 1.42 |
| MolProbity score | 1.78 |
| Clashscore | 6.2 |
| Ramachandran favored / allowed / outliers (%) | 96.8 / 3.2 / 0 |

Extended data collection, refinement and validation statistics are provided in Supplementary Table 1.

An arched helix with the invariant Leu415 at its vertex, shields access to the catalytic iron in P1. The neighboring α2 helix (α2H) distinguishes itself from other LOX homologues. Rather than an elongated helix with six to seven turns, as in 8-LOX and 15-LOX, it is a short three turn helix flanked by extended loops in wildtype 5-LOX^7^. Of note, these two helices are included in three segments, comprising residues Ser172-Ser216, Alα95-Thr302 and Glu418-Gly430, which lack density, likely due to inherent flexibility, thereby preventing reliable modeling. This interpretation agrees well with the fact that the arched helix and α2H were recently suggested to regulate access to the catalytic site via conformational switching between open and closed states^7^. We obtained further support for this notion, as neural network-based classification revealed a class comprising 10 k particles, yielding a low-resolution map with support for α2H in a closed conformation (Supplementary Fig. 2). However, this feature was not maintained when refined to higher resolution.

The catalytic Fe^2+^ is liganded by the terminal carboxylate of Ile674, the amide-oxygen of Asn555 as well as the imidazole groups of residues His368, His373 and His551. The terminal Ile674 is supported by weaker density, compared to the other ligands, extending to cover the entire residue and parts of the preceding Ala673. This suggests increased conformational flexibility at the terminal region of wild type 5-LOX as further explored by MD simulations.

### Architecture of the ATP binding site

In the complex structure, unmodeled density in the vicinity of residues Lys320, Lys656 and Gln657 indicated the position of ATP. The negatively charged phosphates of ATP are held in place by two positively charged residues Lys320 and Lys656. Notably, Lys656 is preceded by Lys655 and Lys654, also recognized as the “KKK” motif of 5-LOX (Fig. 2A, 2B and Supplementary Table 2). The α-phosphate group forms hydrogen bonds with the backbone amide groups of Gln657 and Leu658 (Fig. 2C). Additionally, Lys320 interacts with the γ-phosphate while Gln657 engages the β-phosphate group of ATP. The adenine of ATP is held in place via aromatic interactions with residue Tyr235 and a hydrogen bond between its amino group and the backbone carbonyl oxygens of Leu231 and Ile321 (Fig. 2C). Its ribose is coordinated via its hydroxyl groups by the side-chain amino groups of Lys320 and Lys656 (Fig. 2C). On either side of the ATP molecule, Tyr468 and Tyr235 form π–π interactions in an edge-to-plane geometry, characteristic of many nucleotide-binding proteins, likely guiding the adenine into its correct position. In addition, the aromatic ring of Tyr468 forms a donor-π association with the side chain of Lys320, possibly stabilizing the positioning of Lys320, ribose and γ-phosphate group (Fig. 2C). Along with Tyr235 and Tyr468, other aromatic residues such as Phe470, Tyr471, Tyr660 and Tyr662 appear to participate in aromatic stacking or π–π interactions near the ATP binding pocket and may contribute to structural rearrangements of α-helices and the C-terminal loop upon ATP binding. The binding site of ATP in the cryo-EM structure was also accurately predicted by AlphaFold3^8^.

**Figure 2.**
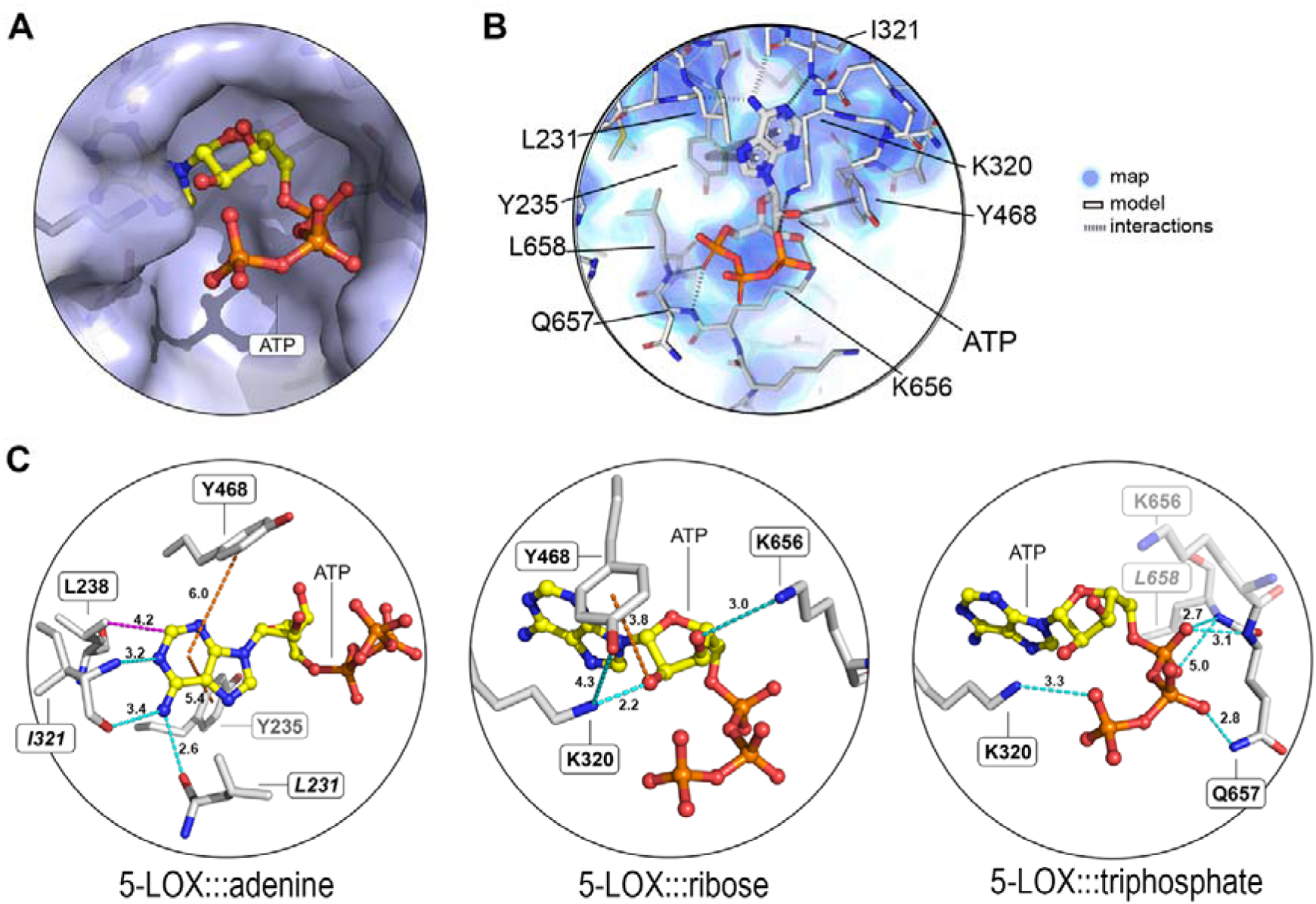
Binding of ATP to 5-LOX. **A** The ATP molecule bound to its cavity. **B** Map and model are shown for ATP and surrounding residues. Atoms of the model are colored as follows: carbon as white or yellow, oxygen as red, nitrogen as blue, sulfur as yellow and phosphorus as orange. **C** Summary of residues or backbone peptide bonds interacting with ATP. Residues involved via backbone amide or carbonyl groups are indicated in italics. Distances of polar, hydrophobic and aromatic interactions are presented with polar (cyan), hydrophobic (purple) and aromatic (orange) dashed lines.

### MD simulations show flexible but stable ATP binding consistent with cryo-EM

MD simulations of the wild-type structure bound to ATP (ATP+), performed in triplicate (total 1,5 µs), showed stable and consistent modes of ATP binding. Across all three simulations, the ATP phosphate groups were predominantly stabilized by Lys320 and Lys656. The amino group of the adenine ring maintained stable hydrogen bond interactions with Ile321 and Leu231 throughout the trajectories, with Leu231 also intermittently contacting the imidazole moiety. Meanwhile, the backbone of Gln657 engaged intermittently with the ribose hydroxyl groups and/or phosphates, whereas its side chain only formed transient contacts in all replicates. The ribose ring was periodically contacted by Lys320, Lys656, Gln657, and Leu658. Finally, Tyr468 established cation–π contacts with the Lys320 or Lys656 side chains in all simulations. These simulations pointed to Lys320, Lys656, and Gln657 as potentially most important in anchoring ATP’s phosphate and ribose moieties, echoing the binding poses observed in the cryo-EM structure.

### ATP spontaneous binding in MD simulations

To evaluate how ATP could spontaneously bind to 5-LOX and to explore potential allosteric effects, we initiated a set of simulations (ATP+out) where ATP was positioned 10 Å from the predicted pocket with its phosphate moiety faced away from the 5-LOX binding site. Overall, these simulations indicate that ATP remains near the binding site for most of each trajectory, primarily through interactions between its phosphate groups and basic residues (Lys320, Lys654, Lys655, and Lys656) located at the pocket entrance. These interactions may facilitate the initial capture of ATP via long-range electrostatic forces. In replicates 1 and 2, ATP stays stably engaged, with replicate 1 even showing spontaneous partial penetration toward the inner cavity—although not to the depth observed in the cryo-EM structure, which would likely require more extensive sampling

### Lys320 is critical for ATP induced allosteric activation of 5-LOX

Selected amino acids with side chains interacting with ATP in the cryo-EM structure were mutated to probe their role in ATP stimulated 5-LOX activity. Exchange of a charged Lys for a hydrophobic Leu at position 320 (Lys320Leu) abolished basal activity (in presence of Ca^2+^ and phosphatidyl choline) and rendered the enzyme insensitive to ATP (Fig. 3A). This appears reasonable in view of this residue’s multiple electrostatic and polar interactions with the nucleotide, observed in the cryo-EM structure (Fig. 2).

**Figure 3.**
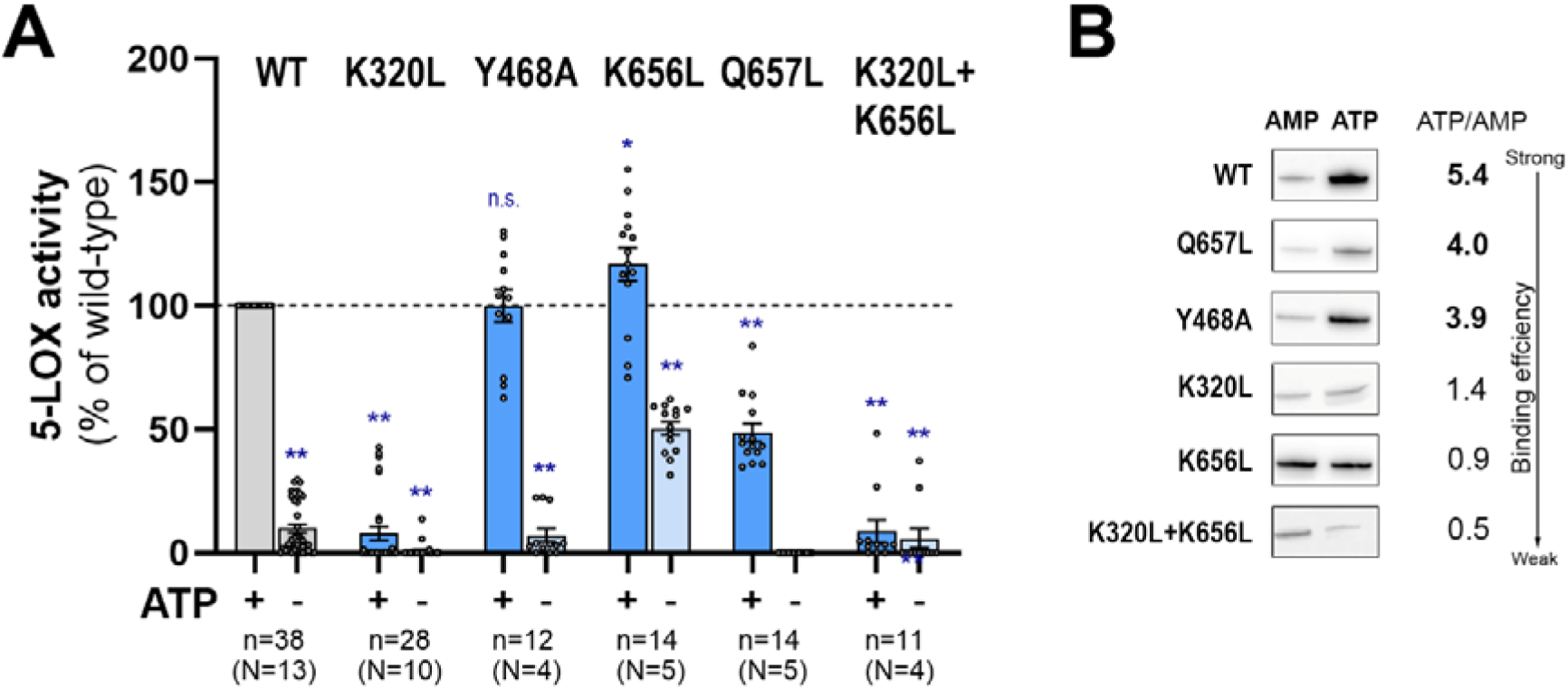
ATP-dependent activation and binding capabilities of wild-type 5-LOX and mutants. **A** Activity assays were performed in the presence of AA (75 µM), Ca^2+^ (0.8 mM), phosphatidylcholine (20 µg/mL), 13S-HpODE (7.5 µM), with (+) or without (−) ATP (3.75 mM) at room temperature for 10 min. Statistical comparisons were made between wild-type 5-LOX (+ATP) and each mutant. All differences are statistically significant (pc<c0.05), except for 5-LOX Y468A, which is indicated as not significant (n.s.). Data are presented as mean ± SEM, based on at least four independent experiments (N). **B** ATP-binding capacity of wild-type 5-LOX (WT) and mutants were assessed by ATP affinity chromatography. Proteins retained on the ATP column were detected by immunoblotting. The ratios of ATP to AMP fractions (ATP/AMP) were determined, reflecting the ATP binding efficiency. See related Supplementary Figures 4 and 5 for panels A and B, respectively.

In contrast, the Lys656Leu mutant had increased 5-LOX activity in both absence and presence of ATP, suggesting that this residue is not critical for ATP induced 5-LOX activation but rather for basal enzyme activity. Interestingly, a similar increase in overall 5-LOX activity, albeit to a lesser extent, was previously reported by Horn *et al.* when mutating Lys656 into the polar Gln^9^ and in fact many other lipoxygenases carry a Leu in this position suggesting that the charged Lys656 may instead play an attenuating role specifically for 5-LOX enzyme activity. This notion is also in line with a previous study suggesting that the three adjacent Lys 654-656, termed the “KKK motif”, in 5-LOX plays a suppressive role for enzyme activity^10^.

When the neighboring Gln657 was exchanged for a Leu, the mutant lost all its basal enzyme activity but responded significantly to ATP, albeit to approx. half of the response exhibited by wild type 5-LOX. Hence, Gln657 does indeed contribute to ATP binding and 5-LOX activation. Removal of the aromatic ring at position 468, yielded the Tyr468Ala mutant whose activity and ATP sensitivity were not significantly different from wild type enzyme. Finally, in the double mutant Lys320Leu/Lys656Leu the Lys320/Leu mutation dominated the phenotype with only slightly improved basal and ATP stimulated activity when combined with the activating Lys656/Leu alteration. Taken together these observations support a crucial role for Lys320 in ATP binding and allosteric activation of 5-LOX. While Gln657 also contributes to the effects of ATP, the role of Lys656 seems to be more important for basal 5-LOX activity than ATP stimulation. Finally, interactions of Tyr468 with ATP do not seem to contribute to allosteric activation of the enzyme.

A qualitative ATP binding assay was performed using ATP-Sepharose affinity columns to assess the relative binding of wild-type 5-LOX and its mutants. Fractions from each purification step were analyzed via Western Blot (Supplementary Fig. 5). The comparison of fractions eluted with ATP or AMP (ATP/AMP) represents the pool of proteins that bind strongly to the column and, after multiple washing steps (including AMP elution), can only be eluted with ATP (Fig. 3B). The relative binding scores (ATP/AMP) demonstrate high ATP affinity for wild-type 5-LOX and the Gln657Leu and Tyr468Ala variants, all of which retained sensitivity to ATP. In contrast, for the Lys320Leu, Lys656Leu and Lys320Leu/Lys656Leu mutants, AMP quite efficiently competed with ATP binding and these mutants showed decreased ATP sensitivity. The seemingly weak binding of Lys656Leu mutant to the ATP-Sepharose suggests that its enhanced enzyme activity is driven by conformational changes rather than ATP binding.

### ATP binding decouples PLAT domain versus catalytic domain motions in 5-LOX

To dissect how ATP modulates 5-LOX dynamics, we performed molecular dynamics simulations of the wild-type enzyme in the absence of ATP (ATP−), with ATP bound (ATP+), or positioned near the binding pocket (ATP+out), totaling nearly 7 μs of sampling with two different force-fields to model iron coordination (see Methods, Supplementary Figs. 6-9). While the first simulation used CHARMM36m with the Won force-field^11^ defaults, well suited to capture large-scale protein motions^12^, the second introduced tighter iron binding parameters, developed by Li-Merz^13^, allowing the comparison of the trade-off between local and global dynamics when modelling ATP binding effects. Our first simulation set revealed rigidity for stable 5-LOX, both global and at the level of the iron coordination site, which remained within canonical distances with virtually absent breathing (see Supplementary Table 3). In contrast, simulations of the wild-type structure (ATP-) revealed increased breathing of the catalytic center, especially upon ATP binding (Supplementary Figs. 10-11, Supplementary Tables 4-5), coupled to increased global flexibility (Supplementary Fig. 12, Supplementary Table 6). In our cryo-EM structure, and in trajectories of wild type enzyme without ATP (ATP-), the PLAT and catalytic domains form a compact interface stabilized by salt bridges linking Arg102 in the PLAT domain to Asp167 and Glu623 in the catalytic domain, together with an Asp80–Arg166 contact. These interactions maintain the enzyme in a closed conformation. However, in ATP+ and ATP+out simulations this contact network is progressively disrupted, allowing up to ~43° rotation of the PLAT domain about the hinge linker (residues 114–125) and increasing the interdomain separation by ~30 Å (Fig. 4A). Root Mean Square Fluctuation (RMSF) profiles and domain center-of-mass distributions confirm this enhanced flexibility under ATP-bound conditions (Fig. 4A, B & Supplementary Fig. 7, 9). Dynamic cross-correlation analysis (DCCM; ATP− minus ATP+) reveals a loss of coordinated motion (Fig. 5C right, red) between the catalytic hinge (residues 114–115) and the flexible C-terminal helix (residues 624–627), consistent with release of interdomain restraints and facilitating large-scale pivoting of the PLAT domain (Fig. 4C; Supplementary Figs. 12-14). These domain-scale changes recapitulate Small-Angle X-ray Scattering (SAXS) observations for the homologous 15-LOX-1 enzyme^14^. Furthermore, residues within the ATP binding pocket show reduced correlation with the PLAT domain in the ATP-bound state (Fig. 5C right, red), while ATP binding induces new or enhanced correlated motions with the catalytic core (Fig. 5C right, blue), reflecting a dynamic rewiring of communication pathways across the enzyme. While simulations performed with the Won Fe^2+^ parameters captured these experimentally reported large-amplitude motions reproducibly, those employing the Li– Merz Fe^2+^ model which enforces a tighter metal coordination with virtually no breathing (Supplementary Fig. 11, Supplementary Table 4) showed only modest oscillations (“trembling”) around the closed state and weaker ATP-dependent transitions (Supplementary Figs. 10 & 11). In this case, ATP only induced a broader span of motion and increased maximal values in some metrics capturing PLAT-catalytic domain distances (See e.g. distance between residues 102-403, Supplementary Fig. 12 & Supplementary Table 6). This difference indicates that the Li–Merz potential enforces a tighter coupling between the catalytic iron and its coordinating ligands, maintaining Fe^2+^-ligand distances near expected values (Supplementary Table 7) and thereby limiting long-range allosteric motions. Importantly, Fe^2+^ remained bound in both the Won (Supplementary Fig. 10) and Li-Merz (Supplementary Fig. 11) models, but in the former, breathing of the catalytic pocket clearly depended on the wild-type sequence and ATP-induction. Of note, quantum chemical calculations of representative structures captured in MD simulations of ATP-, ATP+ and ATP+out, using the Won model, revealed that all Fe^2+^-ligand interactions were overall attractive (negative total energies), indicating that even the most relaxed configurations observed in MD remain energetically stable (Supplementary Fig. 15; Supplementary Table 8).

**Figure 4.**
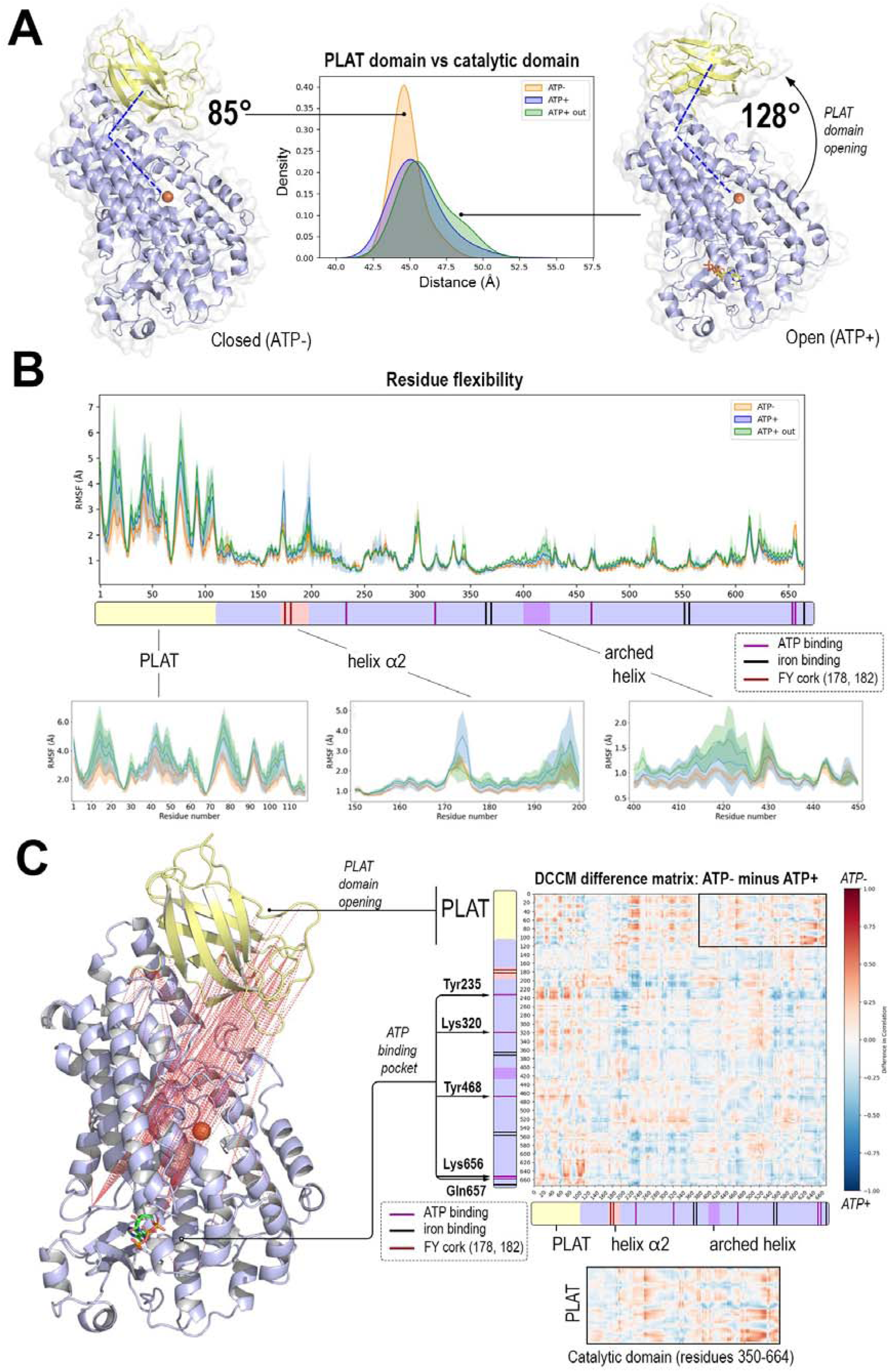
ATP binding allosterically decouples PLAT versus catalytic domain motions in MD simulations of wild type 5-LOX, favoring opening. **A** Distribution of center-of-mass distances between the PLAT and catalytic domains in Won simulations. Representative closed and open states from ATP− and ATP+ trajectories are shown. **B** Root Mean Square Fluctuations (RMSF) across MD simulations of wild type 5-LOX in absence of ATP (ATP−), and with ATP bound (ATP+) or partly bound near its binding site (ATP+out) (3×500 ns each, 1.5 μs per condition). Increased flexibility in the PLAT domain is seen as higher peaks (blue) in RMSF. **C** Right: Dynamic Cross-Correlation Matrix (DCCM) difference map comparing the ATP− and ATP+ states. The heatmap represents correlation differences calculated as DCCM(ATP−) minus DCCM(ATP+). Red regions indicate residue pairs that are more positively correlated in the ATP− state and decoupled in the ATP+ state, while blue regions highlight correlations strengthened in the ATP+ state. White regions represent minimal change between conditions. A representative structural mapping of regions with the strongest correlation differences is shown on the left.

**Figure 5.**
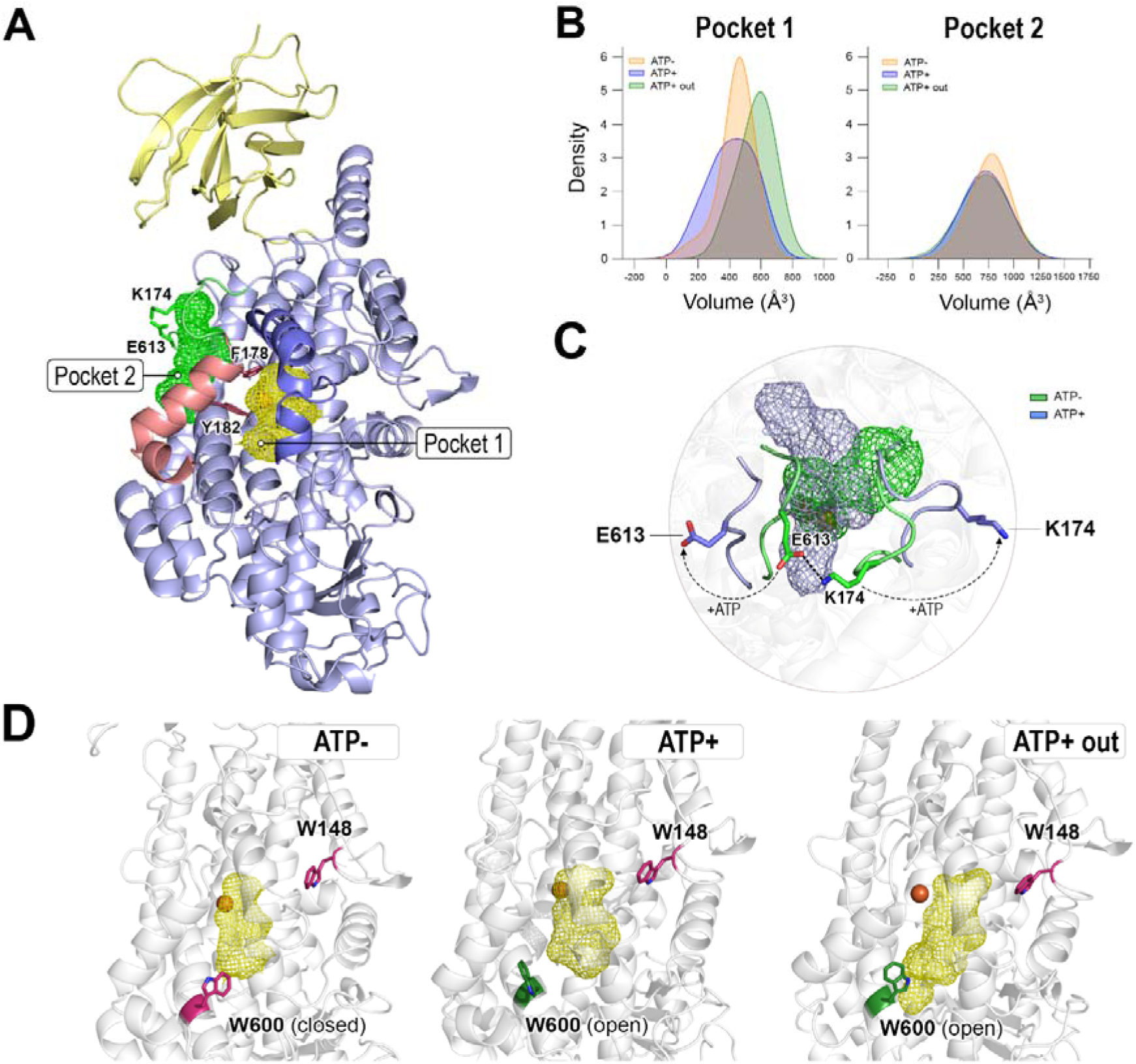
ATP binding induces an increase in the volume of the main catalytic pocket and opening of Trp600. **A** Spatial relationship of the two major pockets detected in MD simulations (snapshot at 120 ns from ATP−). The catalytic pocket (Pocket 1, yellow mesh) extends from the Fe^2+^ center toward the protein surface and is gated by Trp148 and Trp600 at two alternative exits. Pocket 2 (green mesh), is located on the opposite side of the catalytic iron (orange), enclosed at one end by the metal center and at the other by two flexible gate-like loops (green, residues 610–615 and 170–176). For reference, α H is highlighted in pink with the so-called FY-cork represented as sticks (Phe178, Tyr182), while the arched helix is colored in blue. **B** Changes in volume and solvent accessibility are especially significant for Pocket 1 upon ATP interactions (see also Supplementary Figure 16). **C** The gate-like loops covering Pocket 2 exhibit increased opening in the presence of ATP (light blue) and are stabilized in a closed conformation by transient salt bridges in ATP− free simulations (green). Representative snapshots were captured at 465 ns. **D** Trp600 in closed (pink) or open (green) gating in representative snapshots at 190 ns from simulations in absence and presence of ATP. Trp148 (pink) maintains a stable conformation across all simulations. The catalytic pocket (yellow) expands in presence of ATP.

### ATP induced remodeling of cavities and gate loops

In both simulation sets, however, ATP binding also remodels the internal cavity network surrounding the catalytic iron. In Won simulations, concomitant with enhanced fluctuations at the metal center, the catalytic pocket (Pocket 1) in wild type 5-LOX (Fig. 5A, yellow) undergoes pronounced expansion in both volume and solvent-accessible surface area, particularly when ATP adopts the anchored conformation (ATP+out) (Fig. 5B, Supplementary Fig. 16). On the opposite side of the metal center, the second largest pocket detected, Pocket 2 in Fig. 5A (green), is delimited by two flexible, gate-like surface loops (residues 610–615 and 170–176) whose motions are dynamically coupled to the PLAT domain (Fig. 4C & Supplementary Figs. 13-14). These loops open more widely in ATP− bound trajectories, increasing solvent access to the catalytic site. In the ATP− state, they are stabilized in a closed configuration by transient salt bridges (notably Glu613–Lys174) and secondary contacts involving Glu623 and Arg102 from the PLAT domain. Although the PLAT domain does not directly occlude Pocket 2, its closed orientation reinforces this gate through its structural linkage, while ATP binding promotes its relaxation. In addition, the catalytic pocket extends from the Fe^2+^ center toward the surface through two channels capped by Trp148 and Trp600. While Trp148 remains closed, Trp600 alternates between closed and open states and remains predominantly open in ATP− bound trajectories (Fig. 5D), forming a potential “gated” route for substrate entry. Remarkably, in the dynamically restrained Li-Merz simulations, a subtler increase in volume for the catalytic pocket is also seen upon ATP binding (Supplementary Fig. 16). Together, these observations support a model in which ATP acts allosterically to prime 5-LOX for catalysis by coupling nucleotide binding to relaxation of the Fe^2+^ coordination sphere, local cavity breathing and increased PLAT-domain oscillations - motions that may not only facilitate catalysis, but also membrane engagement.

## CONCLUSIONS

The cryo-EM structure of wild-type 5-LOX reveals how ATP engages a positively charged pocket within the catalytic domain to modulate enzyme activity. Functional assays identify Lys320, supported by Gln657, as the key mediator of ATP− dependent activation. Atomistic simulations corroborate these findings and demonstrate how ATP can induce long-range, dynamic conformational changes of the PLAT domain and catalytic center to prime 5-LOX for catalysis and readiness for membrane association. Taken together, our findings offer novel insights to the structural and mechanistic basis of ATP− mediated allosteric regulation in 5-LOX and provide new perspectives for selective modulation of leukotriene biosynthesis.

## MATERIALS AND METHODS

### Materials

The list of materials is presented in Supplementary Table 9.

### Cloning and site-directed mutagenesis

The expression construct containing the codon-optimized Twin-strep-tagged wild-type 5-LOX (TS-wt5-LOX) sequence in the pET21a(+) vector was synthesized by GenScript. The mutations in the wild-type sequence were carried out using Phusion Site-Directed Mutagenesis Kit (Thermo Scientific™). The mutagenesis primers were synthesized by Integrated DNA Technologies. The introduced mutations were validated by sequencing. Plasmids encoding TS-wt5-LOX or mutated TS-5-LOX, which were transformed into *Escherichia coli* BL21(DE3) competent cells.

### Protein preparation

*E. coli* BL21(DE3) cells were grown at 37 °C in in 8 L Terrific Broth medium supplemented with 100 µg/mL ampicillin to an OD600 of 0.5. The protein expression was induced with 0.5 mM IPTG and cells were incubated overnight at 15 °C. Cells were harvested by centrifugation at 5,000 x g for 10 min and suspended in 50 mL of lysis buffer containing 50 mM Tris-HCl (pH 8.0), 150 mM NaCl, 1 tablet of cOmplete™ EDTA-free protease inhibitor cocktail, 1 mg/mL hen egg lysozyme, 250 U of Benzonase® Nuclease. Cells were disrupted by sonication with Sonics Vibra Cell sonicator equipped with the standard tip at 40% amplitude for 1 h (5 s on and 10 s off) and cell debris was removed by centrifugation at 12,000 × g for 1 h at 4 °C. The supernatant was loaded on an immobilized Strep-Tactin® Superflow® column (3 mL; IBA Lifesciences) equilibrated with 100 mM Tris–HCl pH 8, 150 mM NaCl, 1 mM EDTA. The proteins were eluted with 100 mM Tris-HCl pH 8.0, 150 mM NaCl, 1 mM EDTA, 2.5 mM desthiobiotin. The protein was loaded on an 1 mL ATP− agarose column equilibrated with 50 mM Tris-HCl pH 8.0, 150 mM NaCl, 10 mM 2-mercaptoethanol. The column was washed with the same buffer containing 12 mM AMP and eluted with the buffer supplemented with 12 mM ATP. Freshly eluted 0.5 mL fractions were quantified using the Bradford assay and were used immediately for the cryo-EM grid preparation.

### Activity assay

Small-scale expressions of wild-type and mutated 5-LOX were carried out in 10 mL Terrific Broth-ampicillin medium at 15 °C overnight. Cells were harvested in 0.5 mL of 50 mM Tris (pH 8.0) buffer, sonicated with the Sonics Vibra Cell sonicator equipped with a micro tip at 40% amplitude for 5 x 5 seconds and centrifuged at 15,000 x g for 10 min. Next, 50 µL and 3 µL of supernatant was used in incubation assays and Western Blot analysis, respectively. The activity assay was carried in 200 µL of 50 mM Tris (pH 8.0) buffer containing 100 µM arachidonic acid (AA), 150 mM NaCl, 1.2 mM EDTA, 2 mM CaCl_2_, 10 µM 13S-HpODE and 25 µg/mL phosphatidyl choline, in absence or presence of 3.75 mM ATP, for 10 min at room temperature. The reaction was stopped with equal volume of MeOH containing 200 pmol of the 17S-HDoTE standard. 5-LOX products, 5S-H(p)ETE and LTA_4_, were quantified using reverse-phase HPLC. Samples were analyzed on a 3.9 × 150-mm column (C18; Nova-Pak Waters) by eluting products at a flow rate of 1 mL/min with acetonitrile/ water/acetic acid at a ratio of 60:40:0.1 (vol/vol). Absorbance was monitored at 235 nm and the 5-LOX activity was determined in picograms per 1 min in 1 mL, calculated from integrated HPLC peak areas relative to internal standards. The activity of 5-LOX mutants are presented relative to the activity of wild-type 5-LOX and normalized with the protein expression levels determined with Western Blot. Results are presented using the GraphPad Prism program.

### Protein binding assay

Wild-type 5-LOX and mutants were expressed and purified from 200 mL cultures as described in the Protein Preparation section. Aliquots were collected from supernatant, flowthrough, wash and elution steps with 12 mM AMP and 12 mM ATP to assess their binding capacity to the ATP column.

### Western Blot

Proteins were transferred to nitrocellulose membranes using the iBlot2 instrument. The membranes were blocked with 3% milk solution for 1 hour at room temperature. The membranes were treated with in-house 5-LOX primary antibody^15^ at 4 °C overnight followed by incubation with anti-rabbit HRP-conjugated secondary antibody for 1 hour at room temperature. The membranes were developed using SuperSignal™ West Pico PLUS Chemiluminescent Substrate (Thermo Scientific) on the Li-Cor Odyssey instrument.

### Cryo-EM data collection

The grids were glow-discharged for 120 s at 35 mA, then prepared in the following manner: 3.5 µL of sample at 50 µg/ml were applied two times on UltrAuFoil 1.2/1.3 300 mesh Au grids and plunge-frozen in liquid nitrogen-cooled liquid ethane using a Vitrobot Mark IV (Thermo Fisher Scientific) with a blot force of 3 and 5 s blot time, at 4 °C and 100% humidity. Grids were clipped and loaded into a 300 kV Titan Krios G3 electron microscope with cold field emission gun (Thermo Fisher Scientific, EPU software). Grids were screened for quality based on particle distribution and density, and images from the best grid were recorded. Data were acquired using a Gatan K3 direct electron detector with Bioquantum energy filter (10 eV) in counting mode at a nominal magnification of 165 000 x, corresponding to a pixel size of 0.5076 Å. A total of 8600 movies were recorded with an accumulated electron dose of ~65 e−/Å². Details of the other data collection parameters used for each sample are given in Supplementary Table 1.

### Cryo-EM SPA processing and model building

8205 movies were pre-processed applying batch motion- and CTF-correction jobs in cryoSPARC^16^. 993 micrographs were discarded interactively by thresholding the CTF fit resolution to below 10 Å, the average defocus to below 20,285 Å and the relative ice thickness to between 0.7 and 1.3. Blob picking with minimal and maximal diameters of 80 and 200cÅ, respectively, followed by particle inspection yielded 1,607,961 particles. These particles were extracted with a box size of 400 binned to 128 pixels, corresponding to a pixel size of 1.586 Å. 357,451 particles were selected in 37 out of 200 2D classes (Supplementary Fig. 2A), aligned in three final full and 50 online EM iterations with a batch size of 200, a maximum resolution of 8 Å and a circular mask of 120 Å. Four volumes comprising 24 079, 40 493, 15 172 and 20 256 particles were reconstructed *ab initio* from a subset comprising 100 k particles from the selected 2D classes. Iterative heterogeneous refinements with four volumes were applied to isolate a stack of 130,978 particles that was refined homogeneously to a resolution of 3.3 Å. After re-extracting the particles with a box size of 400 binned to 192 pixels (1.0575 Å pixel size), non-uniform refinement 2 improved the resolution of the reconstruction to 3.05 Å. However, the map exhibited streaky features and conical FSC (cFSC) summary plots revealed a wide directional resolution range between 6.79 and 2.75 Å. Furthermore, the cFSC Area Ratio (cFAR) value of 0.08 indicated strong orientation bias (Supplementary Fig. 2B and 2C). After importing the particle stack into Relion with the help of the UCSF pyem suite^17, 18^, 3D auto-refinement with Blush regularization (Relion-blush)^19^ led to a significantly improved reconstruction (Supplementary Fig. 2B and 2C). The obtained map appeared more isotropic, the cFSC resolution range became narrower (3.92 → 2.72 Å) and the cFAR metric increased to 0.31. The particle stack was downsampled to 64 pixels (3.1725 Å pixel size) and analyzed in CryoDRGN^20, 21^ by training the neural network with an 8-dimensional latent variable model for 50 epochs on the default 1024×3 architecture. Iterative conformational landscape analysis with ward linkage for agglomerative clustering resulted in three classes comprising 97,636, 23,181 and 9,618 particles. 3D auto-refinement using Relion-blush of each class revealed a significantly worse reconstruction for the 97 k class (Supplementary Fig. 2C) compared to the other two classes. While the initial low resolution map of the 10 k particle stack indicated map support for the a2H in a closed conformation (Supplementary Fig. 1C), this feature was not maintained when refined to higher resolution. The two classes comprising 23 k and 10 k particles were combined for the final Relion-blush 3D auto-refinement (Supplementary Fig. 2B). The narrower cFSC resolution range (3.61 → 2.95 Å) and increased cFAR metric (0.56) of the high quality reconstruction indicated further map improvement. The refined wild-type 5-LOX structure exhibits Molprobity evaluation score and model-to-map cross-correlation (cc) values of 1.24 and 0.8, respectively, demonstrating a physically valid model with excellent map support for 90% of the model. The primary and secondary maps deposited under EMDB code 52771 were post-processed applying Phenix’s model-based local anisotropic sharpening^22^ with an effective average B-sharpening value of −53 Å^2^ and EMready^23^ with the deposited mask, respectively. The recommended viewing threshold levels of the primary, secondary and half-maps are at values of 5, 0.01 and 0.01, respectively. The AlphaFold-predicted^24, 25^ structure of 5-LOX [https://alphafold.ebi.ac.uk/entry/P09917] was docked into the map using Phenix^26^. The catalytic iron was transferred from the superimposed crystal structure of the stable 5-LOX mutant [https://www.rcsb.org/structure/3o8y]^3^. The position of the ATP ligand was predicted accurately by AlphaFold3^8^ to occupy an unmodeled density located in the vicinity of residues K320 and Q657. The model was refined and validated iteratively using Coot, Isolde, Phenix’s real space refinement and Molprobity^27, 28, 29, 30^. The structure was analyzed and visualized using PyMOL^31^, ChimeraX, Arpeggio, VMD and custom-made scripts in Python and Rstudio^32, 33, 34, 35, 36, 37^.

### Cavity mapping and molecular docking

Binding pockets and cavity volumes in 5-LOX were calculated using the PyVOL tool (v1.7.6) within the PyMOL molecular visualization program. Pocket detection was performed with a minimum radius of 1.5 Å (1.2 Å for narrower cavities), a maximum probe radius of 5 Å, and a minimum pocket volume threshold of 200 Å³. Interaction sites between the catalytic pocket of 5-LOX and AA or 5S-HpETE were predicted via molecular docking using the cryo-EM structure of 5-LOX as a template. Ligands were prepared in Avogadro (v1.2.0). Prior to the docking, all non-essential molecules, except the catalytic non-heme iron, were removed from the cryo-EM structure. Simulations were carried out with AutoDock Vina (v1.2) through the UCSF Chimera (v1.19) interface, with the exhaustiveness parameter set to 8. Binding modes with the best docking scores (kcal/mol) were visualized using PyMOL (v3.1.5.1).

### Molecular dynamics simulations and analysis

MD simulations were performed using the cryo-EM structure of wild-type human 5-LOX in three distinct conditions: (1) ATP−, in which ATP was removed and the structure relaxed; (2) ATP+, with ATP bound in the nucleotide pocket; (3) ATP+ out, where ATP was displaced 10 Å from the pocket with its phosphate moiety facing away to prevent immediate interactions. This design enabled exploration of both bound and capture-like ATP states. Each system was prepared independently using CHARMM-GUI, which imported the experimental coordinates, built the simulation box, added solvent and ions, and applied periodic boundary conditions. Protein parameters were described by the CHARMM36m force field^38^, while ATP parameters were taken from the CHARMM36 all-hydrogen nucleic-acid topology. The Fe^2+^ cofactor was represented using the CHARMM ion parameters developed by Won (Won *et al.* 2012)^11^, which accurately captured the stability of the Fe binding site in preliminary simulations of stable 5-LOX (see below), and re-run again with tighter Fe^2+^ parameters by Li-Merz (Li *et al.* 2013)^13^ obtaining similar results. Each system was solvated in a rectangular TIP3P water box^39^ extending 10 Å from the solute, and neutralized with K^+^ and Cl^-^ ions at 0.15 M using Monte Carlo ion placement. All titratable residues were assigned standard protonation states at pH 7.0 according to CHARMM-GUI defaults, including appropriate histidine tautomers. All simulations were carried out with GROMACS 2024.3^40^ using periodic boundary conditions. Each system underwent energy minimization by steepest descent (converged at Fmax ≤ 1000 kJ mol^-1^ nm^-1^), followed by equilibration in the NVT ensemble (constant number of particles, volume, and temperature) for 125 ps with a 1 fs timestep. Production simulations were then performed in the NPT ensemble (constant number of particles, pressure and temperature) at 303.15 K with three independent replicas per system, each initiated with randomized Maxwell–Boltzmann velocities. Stable 5-LOX was simulated for 600 ns in two replicas, while for the ATP−, ATP+, and ATP+out systems, each replicate was run for 500 ns (2 fs timestep; 2.5 x 10^8^ steps). The velocity-rescale thermostat and Parrinello–Rahman barostat were used for temperature and pressure coupling, respectively. Long-range electrostatics were treated with particle-mesh Ewald, and covalent bonds to hydrogen were constrained using LINCS.

Trajectory post-processing was performed with GROMACS utilities. Each trajectory was centered on the protein, periodicity was removed, and frames were extracted every 1 ns. For subsequent analyses, three replica trajectories were concatenated per condition to perform principal component analysis (PCA) and define global motions in a unified structural space. Analyses included RMSD, RMSF, domain distance measurements, iron-coordination dynamics, and dynamic cross-correlation matrices (DCCM) to quantify allosteric coupling. All analyses were conducted using MDAnalysis (v2.7.0)^41, 42^ with Pandas^43^ and NumPy^44^, and Matplotlib and Seaborn^45^ for statistical treatment and visualization. Cavities were identified with MDpocket from the Fpocket suite^46^, and molecular graphics were generated in PyMOL^31^.

## Acknowledgments

We are grateful to Dr. Martin Hällberg and the Karolinska Institutet 3D-EM facility: https://ki.se/cmb/3d-em and computational resources from the Swedish National Infrastructure for Computing (NAISS 2023/5-400, NAISS 2024/5-88, NAISS 2024/1-7 and NAISS 2024/3-5)

## Author contributions

T.T. produced 5-LOX, carried out mutational analysis, and collected cryo-EM data. MD simulations were carried out and analysed by F.P. and L.O.. A.N.H. carried out quantum chemical calculations O.R. helped analyze data, gave intellectual input and revised the MS. T.T. did initial data processing and T.S. solved the cryo-EM structure. T.T, F.P., L.O., T.S. and J.Z.H wrote the manuscript J.Z.H conceptualized, planned and supervised the project.

