## Supplementary Information for "Structural basis for ATP regulation of human 5-lipoxygenase"

**Supplementary results**

Supplementary Figures 1-16

Supplementary Tables 1-8

**Matrials and Methods**

Supplementary Tables 9-11

Supplementary Methods 1

**RESULTS**

**Supplementary Figure 1.**

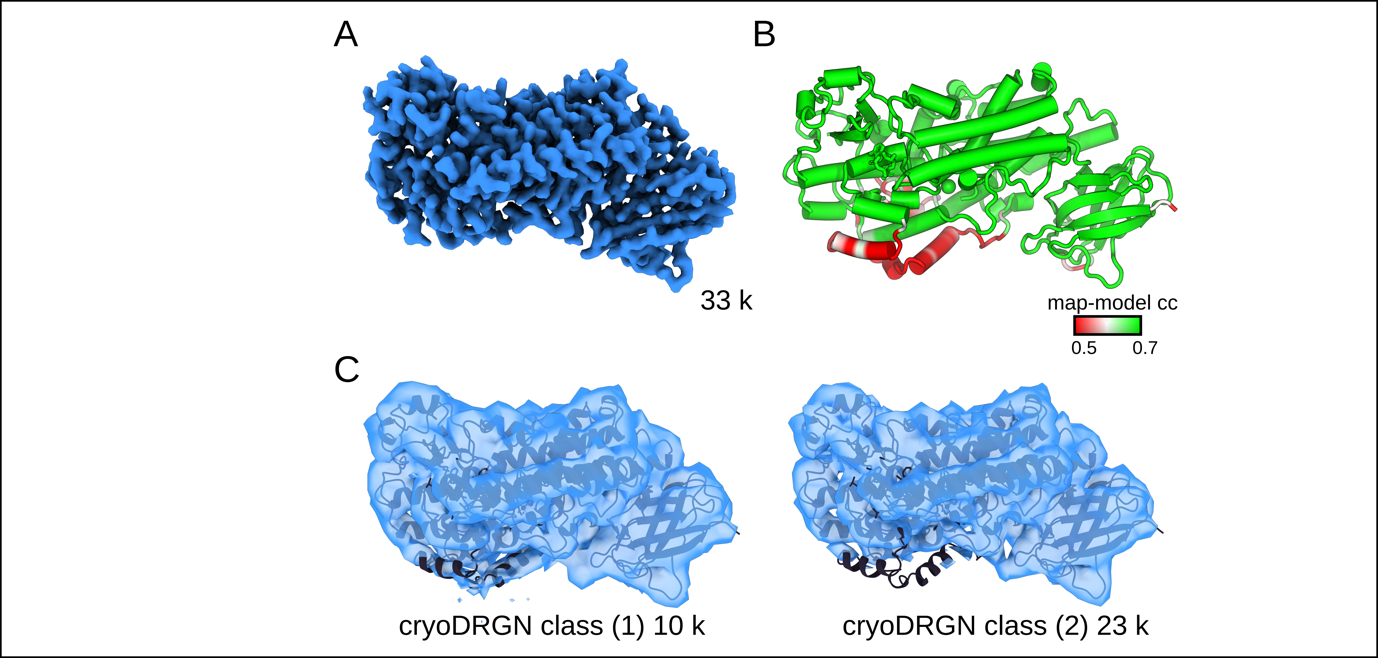

**Map-model correlation and cryoDRGN classification related to Figure 1.**  **A** The 3.3 Å cryo-EM map of 5-LOX is shown in blue at a map level of 6.0σ. **B** The model of 5-LOX is colored in red, white and green for map-model cross correlation values of 0.5, 0.6 and 0.7, respectively, corresponding to weak, middle and strong map support. **C** Neural network-based classification using ‘cryoDRGN’ revealed two classes comprising 10 k and 23 k particles. The two classes were combined for the final reconstruction shown in panel A. The low resolution map of class 1 exhibits density for the α2 helix in a closed conformation. This feature was not maintained when refined to higher resolution. The model and maps are shown as black cartoon and semi-transparent blue surfaces, respectively.

**Supplementary Figure 2.**

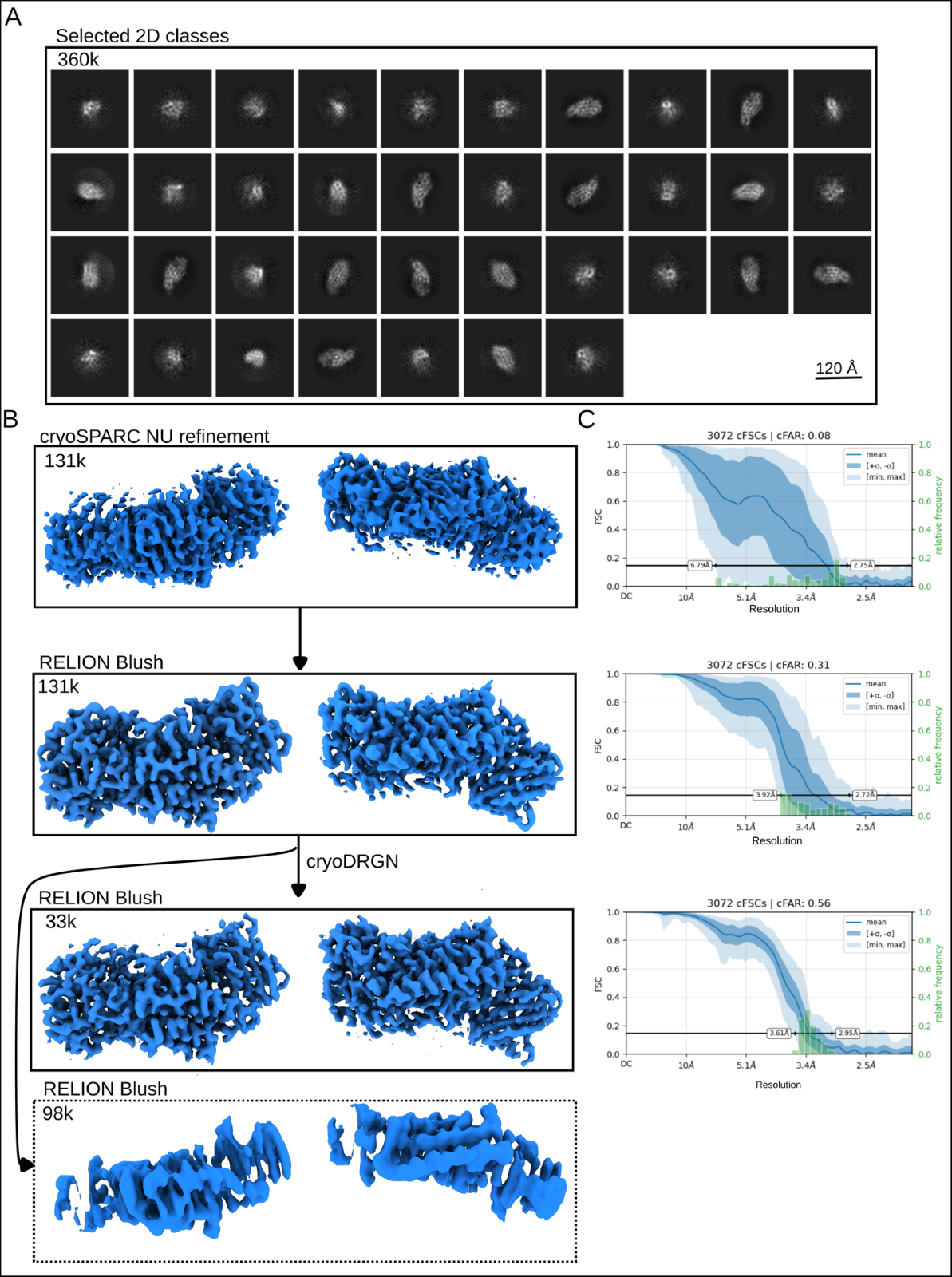

**2D classes and critical processing steps.**  **A** The depicted 2D classes comprising 360k particles were selected for further analysis. **B** Maps of critical SPA processing steps. The application of Blush regularization to the reconstruction has significantly improved map appearance. Conformational landscape analysis in cryoDRGN identified a stack of 33k particles yielding a map virtually identical to the reconstruction derived from 131k particles. More details are described in the main text. **C** Conical FSC (cFSC) summary plots document the increased directional signal content at later stages of the processing pipeline. cFSC summary plots are derived from a set of cFSC curves with conical axes sampled along a uniform spherical distribution. The blue line and shades visualize the mean, standard deviation, minimum and maximum values of the correlations at each spatial frequency. The histogram of 0.143 crossings, corresponding to the spread of resolution values over direction, is overlaid in green. The cFSC Area Ratio (cFAR) metric ranges between 0 and 1 for strong and missing orientation bias, respectively.

**Supplementary Table 1.**

| **Data collection** |  |
| --- | --- |
| Magnification | 165 000x |
| FFVoltage (kV) | 300 |
| Electron exposure (e^-^/Å^2^) | 65 |
| Defocus range (µm) | –0.8 to –2.5 |
| Pixel size (Å) | 0.5076 |
| Number of movies |  |
| collected | 8 205 |
| curated | 7 212 |
| **Data processing** |  |
| Number of Particles |  |
| cs Blob picked (curated) | 1 607 961 |
| cs 2D class selected | 357 451 |
| cs NU refined | 130 978 |
| Relion final stack | 32 799 |
| **Model** |  |
| Composition (#) |  |
| Chains | 2 |
| Atoms | 5533 (Hydrogens: 0) |
| Residues | Protein: 674  Nucleotide: 0 |
| Water | 0 |
| Ligands | FE2: 1 |
|  | ATP: 1 |
| Bonds (RMSD) |  |
| Length (Å) (# > 4sigma) | 0.003 (0) |
| Angles (°) (# > 4sigma) | 0.692 (0) |
| MolProbity score | 1.38 |
| Clash score | 4.57 |
| Ramachandran plot (%) |  |
| Outliers | 0.00 |
| Allowed | 2.83 |
| Favored | 97.17 |
| Rama-Z (Ramachandran plot Z-score, RMSD) |  |
| whole (N = 672) | -0.82 (0.30) |
| helix (N = 281) | 0.08 (0.28) |
| sheet (N = 75) | -0.05 (0.58) |
| loop (N = 316) | -1.19 (0.33) |
| Rotamer outliers (%) | 0.00 |
| Cbeta outliers (%) | NA |
| Peptide plane (%) |  |
| Cis proline/general | 0.0/0.0 |
| Twisted proline/general | 0.0/0.0 |
| CaBLAM outliers (%) | 0.75 |
| ADP (B-factors) |  |
| Iso/Aniso (#) | 5533/0 |
| min/max/mean |  |
| Protein | 76.25/278.27/139.48 |
| Nucleotide | --- |
| Ligand | 104.97/162.83/126.39 |
| Water | --- |
| Occupancy |  |
| Mean | 1.00 |
| occ = 1 (%) | 100.00 |
| 0 < occ < 1 (%) | 0.00 |
| occ > 1 (%) | 0.00 |
| **Data** |  |
| Box |  |
| Lengths (Å) | 69.80, 68.74, 115.27 |
| Angles (°) | 90.00, 90.00, 90.00 |
| Map pixel size (Å) | 1.0575 |
| Supplied Resolution (Å) | 3.0 |
| Resolution Estimates (Å) | Masked Unmasked |
| d FSC (half maps; 0.143) | 3.2 3.4 |
| d 99 (full/half1/half2) | 3.5/2.1/2.1 3.2/2.1/2.1 |
| d model | 3.4 3.4 |
| d FSC model (0/0.143/0.5) | 2.2/2.7/3.4 2.4/2.9/3.5 |
| Map min/max/mean | -0.01/0.02/0.00 |
| **Model vs. Data** |  |
| CC (mask) | 0.81 |
| CC (box) | 0.78 |
| CC (peaks) | 0.71 |
| CC (volume) | 0.81 |
| Mean CC for ligands | 0.84 |

**Cryo-EM data collection and refinement statistics**

**Supplementary Table 2.**

| **Residue** | **Interacting atom / bond** | **ATP Moiety** | **Interaction** | **Distance (Å)** |
| --- | --- | --- | --- | --- |
| **Tyr235** | Side chain | Adenine (6C ring) | π–π edge | 5.4 |
| **Tyr235** | Side chain | Ribose C5' | Aliphatic | 3.4 |
| **Tyr468** | Side chain | Adenine (6C ring) | π–π edge | 6.0 |
| **Tyr468** | Side chain | Ribose C2'-OH | Donor–π | 3.8 |
| **Leu231** | Backbone carbonyl | Adenine C6-NH2 | H-bond (acceptor) | 2.6 |
| **Leu238** | Side chain | Adenine C2 | Hydrophobic | 4.2 |
| **Ile321** | Backbone amide | Adenine N1 | H-bond (donor) | 3.2 |
| **Ile321** | Backbone carbonyl | Adenine C6-NH2 | H-bond (acceptor) | 3.4 |
| **Lys320** | Side chain | Ribose C2'-OH | H-bond | 2.2 |
| **Lys320** | Side chain | γ-phosphate | Ionic | 3.3 |
| **Lys320** | Side chain | Tyr468 OH | H-bond | 4.3 |
| **Lys320** | Side chain | Adenine | Aliphatic | 4.1 |
| **Lys656** | Side chain | Ribose C3'-OH | H-bond (donor) | 3.0 |
| **Lys656** | Side chain | Ribose C5' | Aliphatic | 3.9 |
| **Gln657** | Side chain | β-phosphate | Ionic | 2.8 |
| **Gln657** | Backbone amide | α-phosphate oxygen | H-bond (donor) | 3.1 |
| **Leu658** | Backbone amide | α-phosphate oxygen | H-bond (donor) | 2.7 |
| **Leu658** | Side chain | Ribose C5' | Hydrophobic | 4.3 |

**Interactions and distances within the ATP binding pocket of 5-LOX.**

**Supplementary Figure 3.**

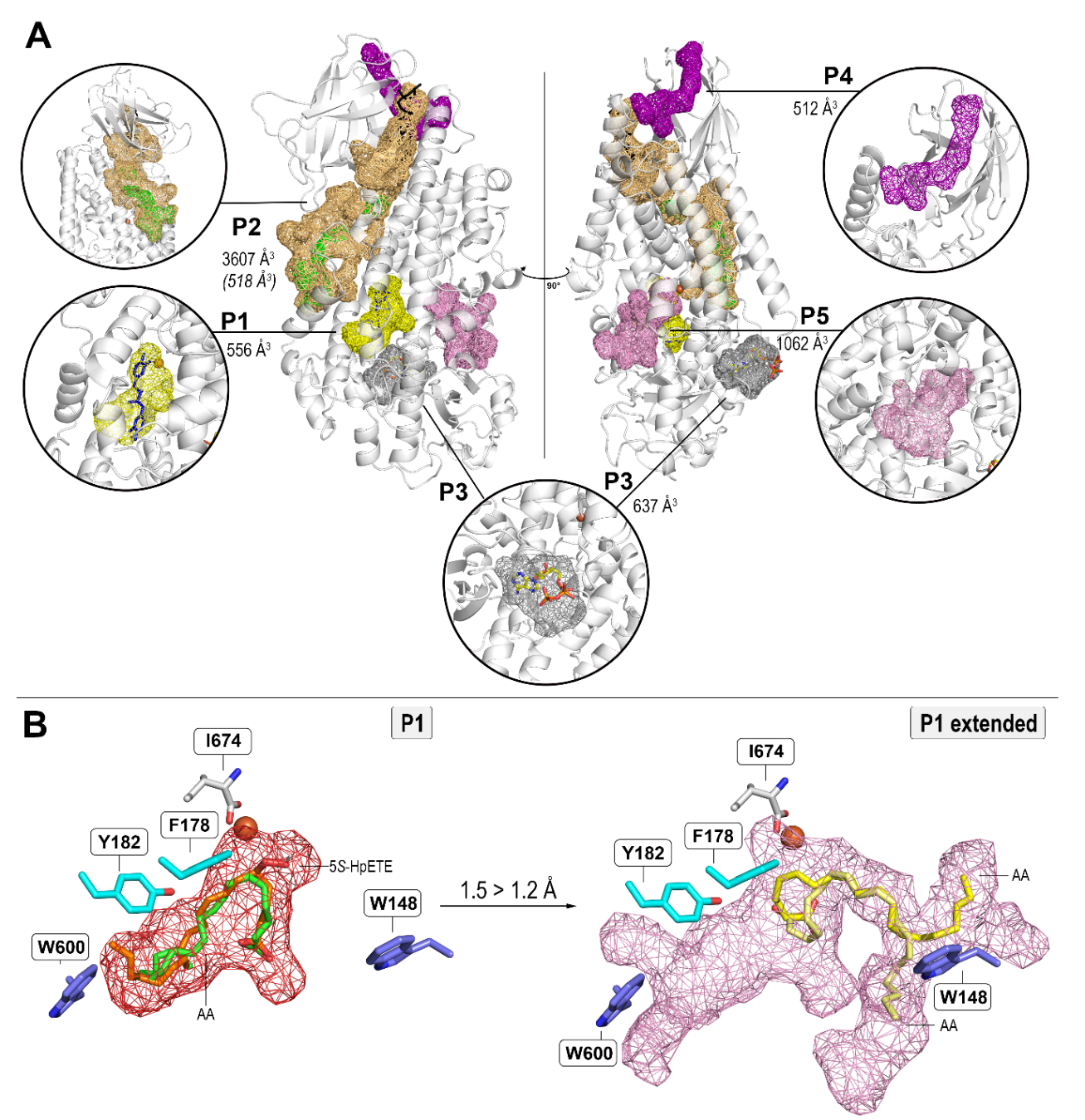

**Measured cavities of wild-type 5-LOX.** **A** Five major cavities (P1–P5) were identified using a probe radius range of 1.5-5.0 Å and a minimum cavity volume of 200 Å³. The cavity volumes are presented in Å^3^ and displayed as volumetric meshes and color-coded as follows: P1 (yellow), P2 (orange), P3 (grey), P4 (purple) and P5 (pink). P1 corresponds to the proposed catalytic site for fatty acid substrates. P2 forms part of the iron-coordinating center and extends toward the interface between the catalytic and PLAT domains (not shown in Fig. 1E). Within P2, a sub-pocket (green) reflects the cavity space located exclusively in the catalytic domain (see Fig. 1E,F), excluding the PLAT contribution. P3 represents the ATP binding pocket. P4 is located in the hinge region connecting the catalytic and PLAT domains. P5 is positioned on the opposite side of the catalytic site. The AKBA inhibitor, located between the catalytic and PLAT domains within P2, is shown in black. The NDGA inhibitor occupying P1 is shown in blue. **B** Docking of AA (green) and 5S-HpETE (orange) into the P1 cavity and AA (yellow) into the extended P1 cavity with a 1.2 Å opening. The cavity mesh is shown in red (P1) and pink (extended P1). W600 and W148, acting as lids of the substrate channel, and the FY cork are presented in purple and cyan, respectively. The catalytic iron, coordinated by the C-terminal isoleucine (white), is depicted as an orange sphere.

**Supplementary Figure 4.**

| 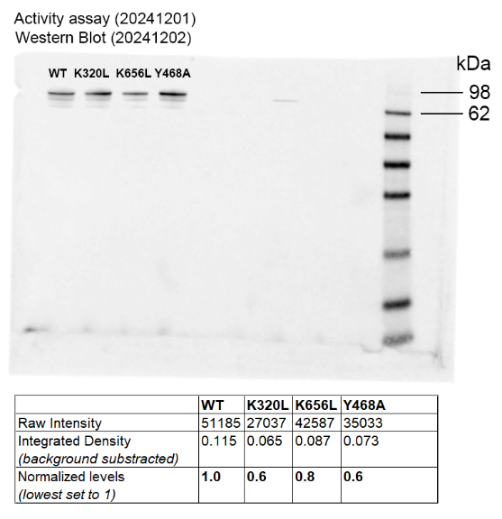 | 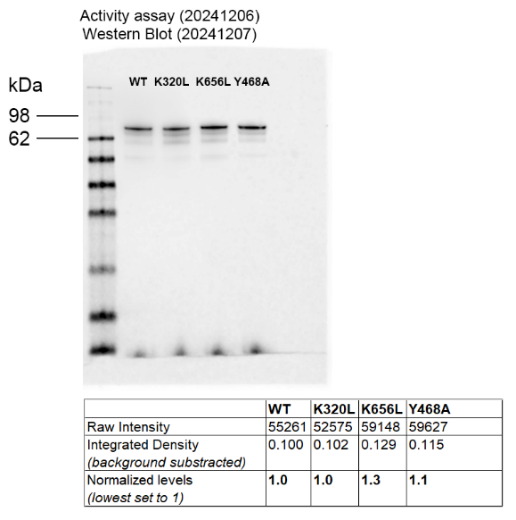 |
| --- | --- |
| 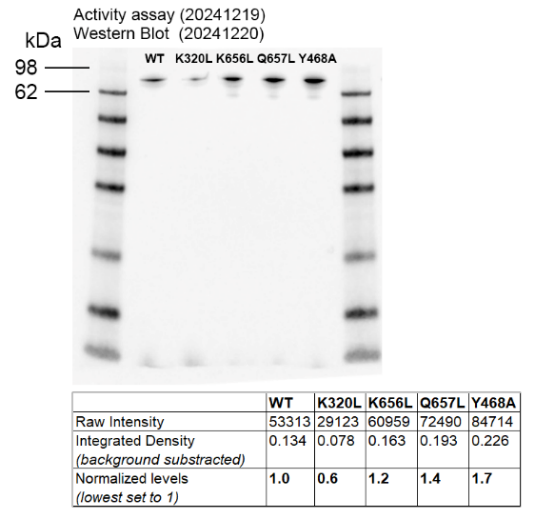 | 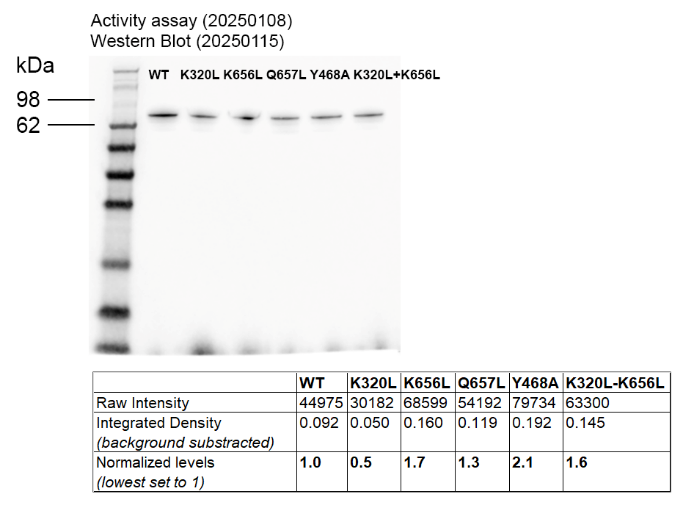 |
| 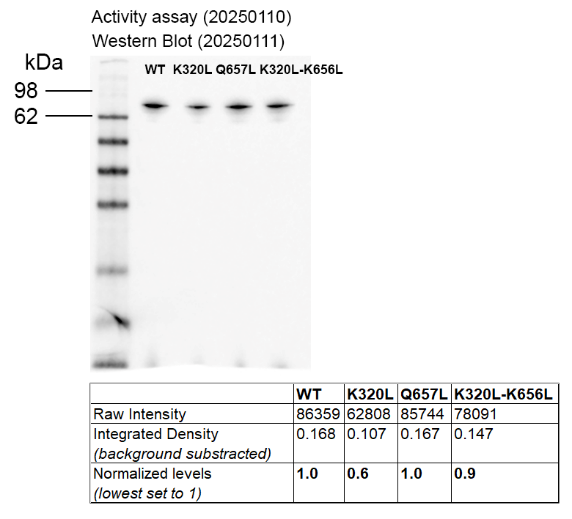 | 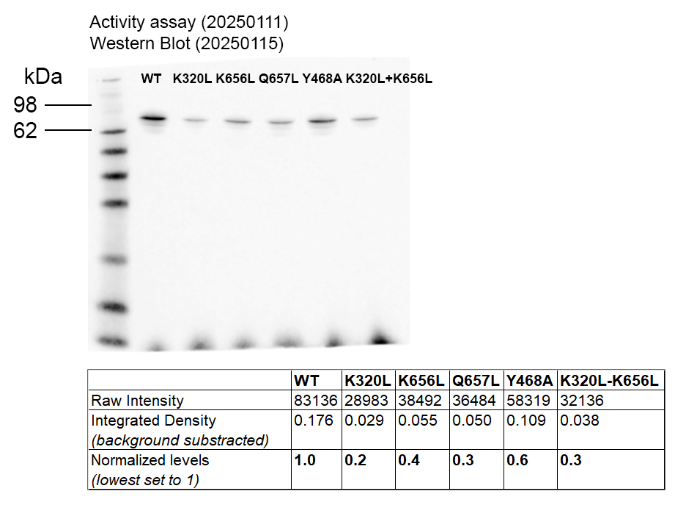 |
| 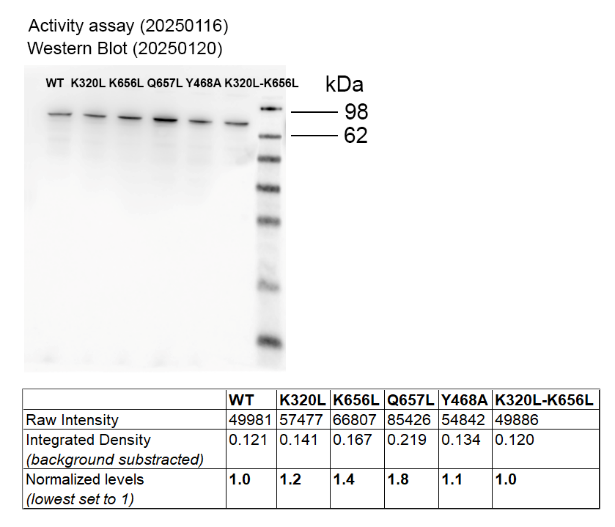 | 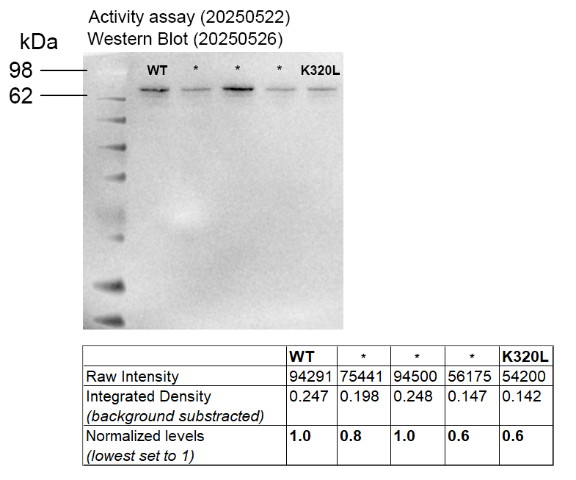 |
| 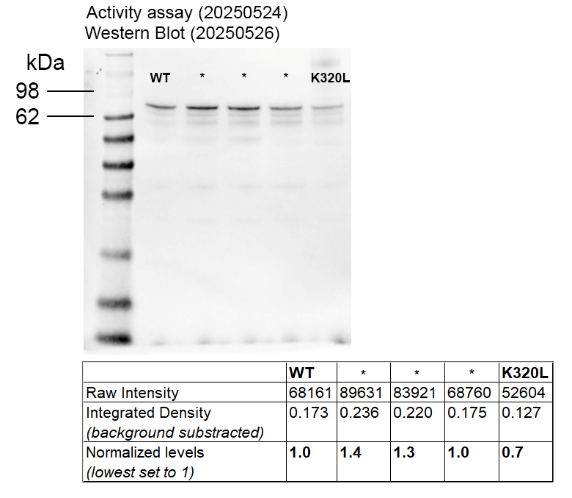 | 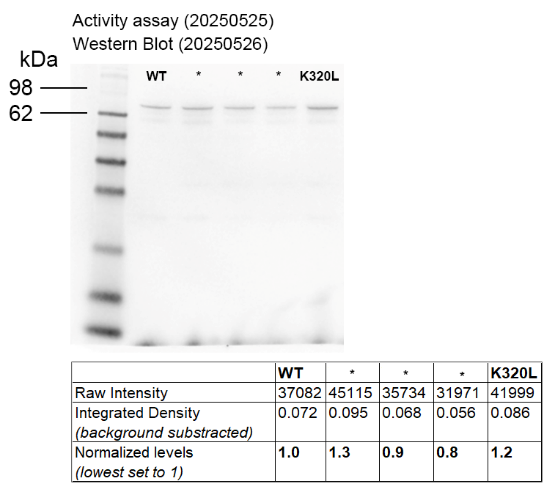 |
| 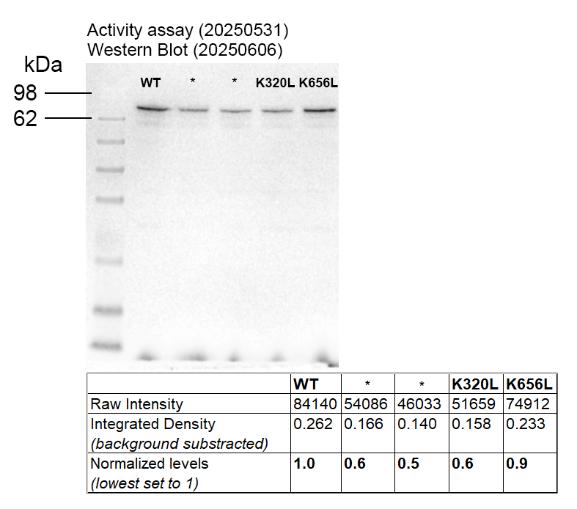 |  |

**Quantification of band intensities and normalization of protein levels.** Protein band intensities were quantified using ImageJ based on integrated density values. Ratios were calculated relative to wild-type 5-LOX and used to normalize enzymatic activity measurements. * - these mutations were not relevant to this study.

**Supplementary Figure 5.**

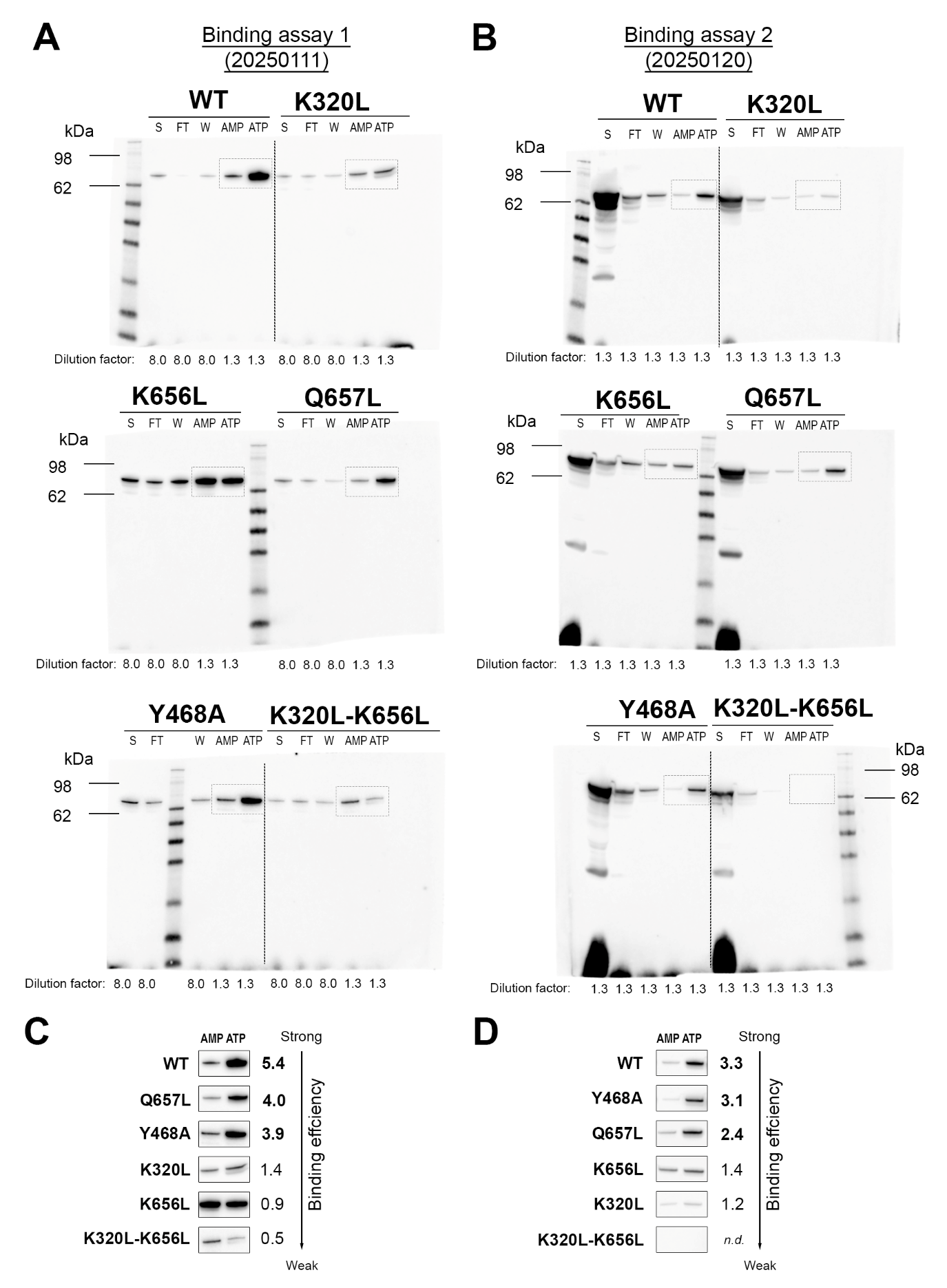

**ATP binding capacity of wild-type 5-LOX (WT) and mutants. A,B** ATP binding capacity of wild-type 5-LOX (WT) and the indicated mutants was assessed by ATP–sepharose affinity chromatography. Proteins retained on the ATP-sepharose column from two independent binding assays (20250111 and 2025012) were detected by immunoblotting. The supernatant (S) of cell lysate and flow-through (FT) samples were diluted 8-fold in 20250111, whereas the wash (W), the adenosine monophosphate (AMP) and adenosine triphosphate (ATP) eluates were diluted 1.3-fold. In the 20250120 binding assay, only 1.3-fold dilutions were carried out. **C,D** Densitometric quantification of band intensities from A and B respectively, determined with ImageJ. The binding scores were calculated considering density and represent the ratio of ATP to AMP fractions (ATP/AMP), reflecting the ATP binding efficiency. Ratios ≥ 2 and 0.5–2 indicate strong and weak binding, respectively.

**Supplementary Figure 6.**

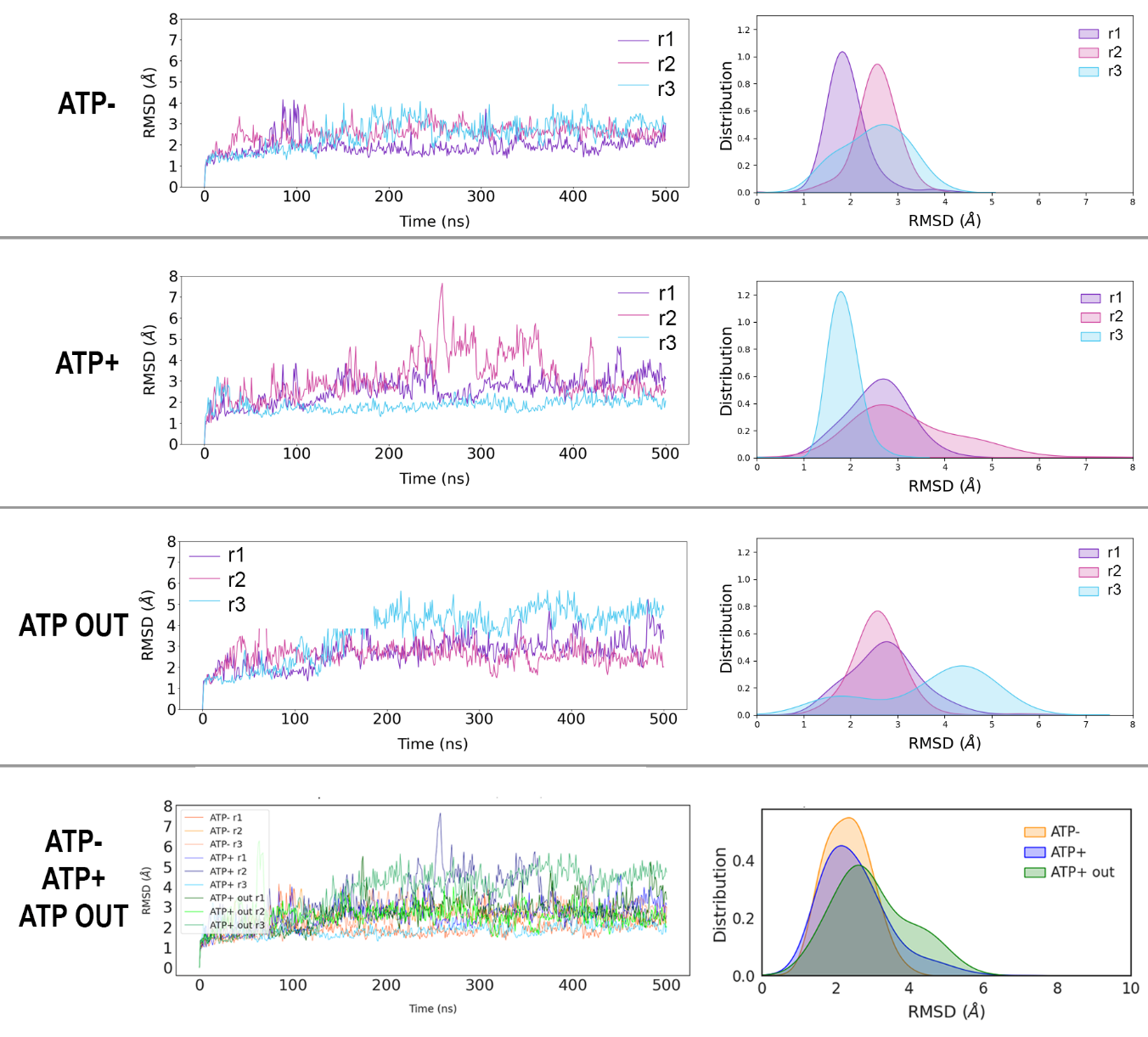

**Root-Mean-Square-Deviations (RMSD) versus the starting structure in CHARMM36m-Won simulations^1^.**

**Supplementary Figure 7.**

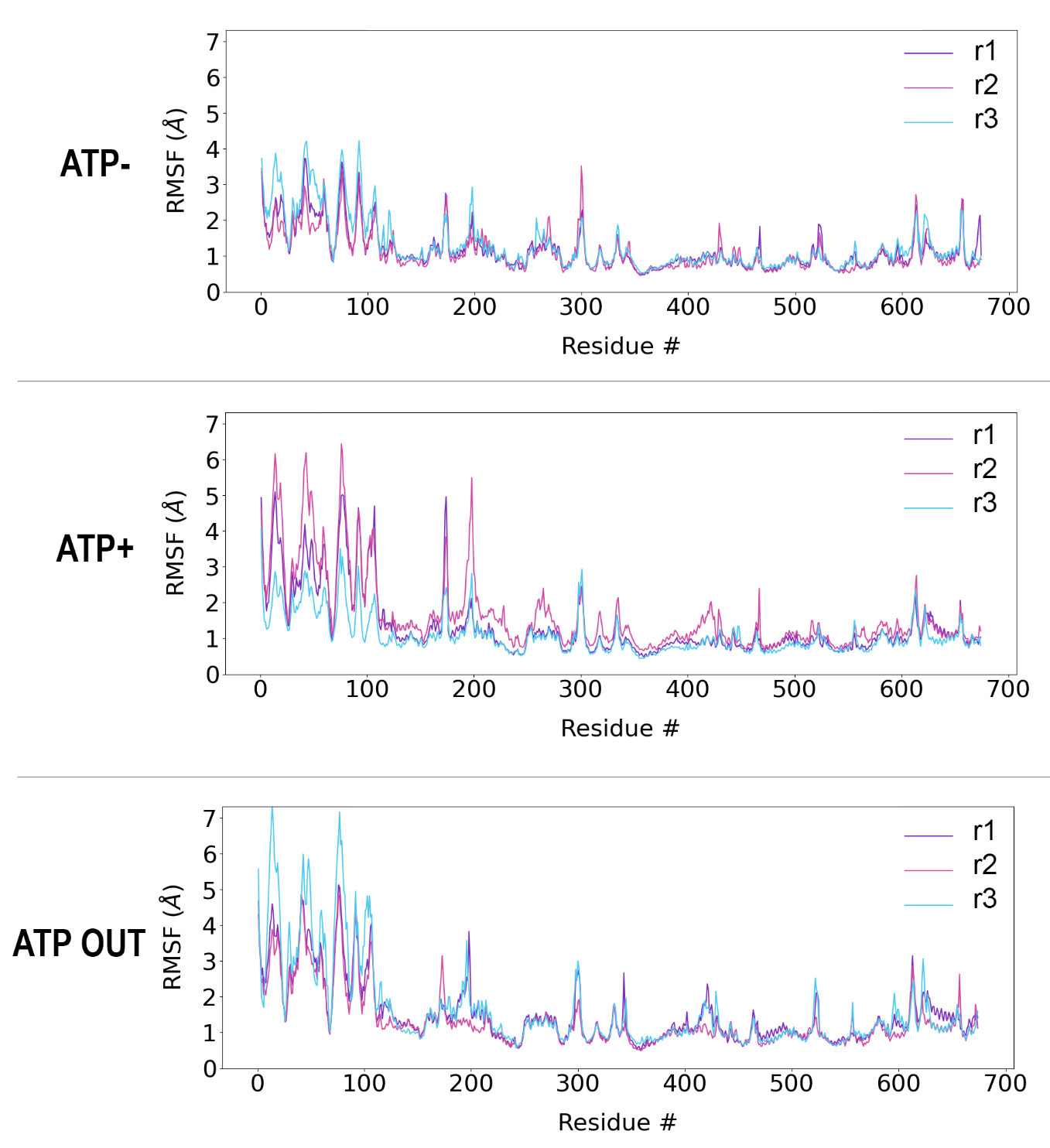

**Root-Mean-Square-Fluctuations (RMSF) per residue in CHARMM36m-Won simulations^1^.**

**Supplementary Figure 8.**

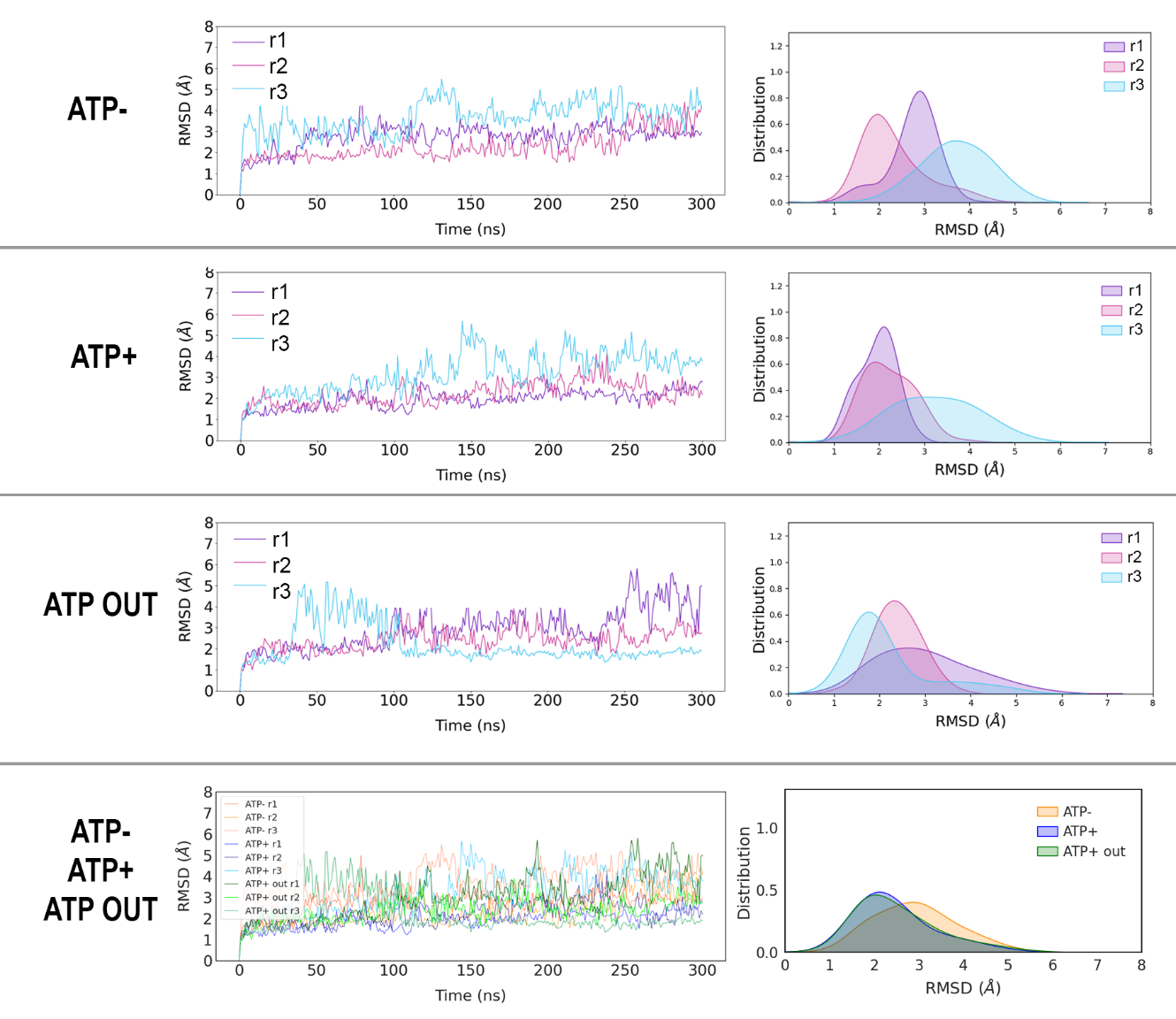

**Root-Mean-Square-Deviations (RMSD) versus the starting structure in CHARMM36m-Li-Merz simulations^2^.**

**Supplementary Figure 9.**

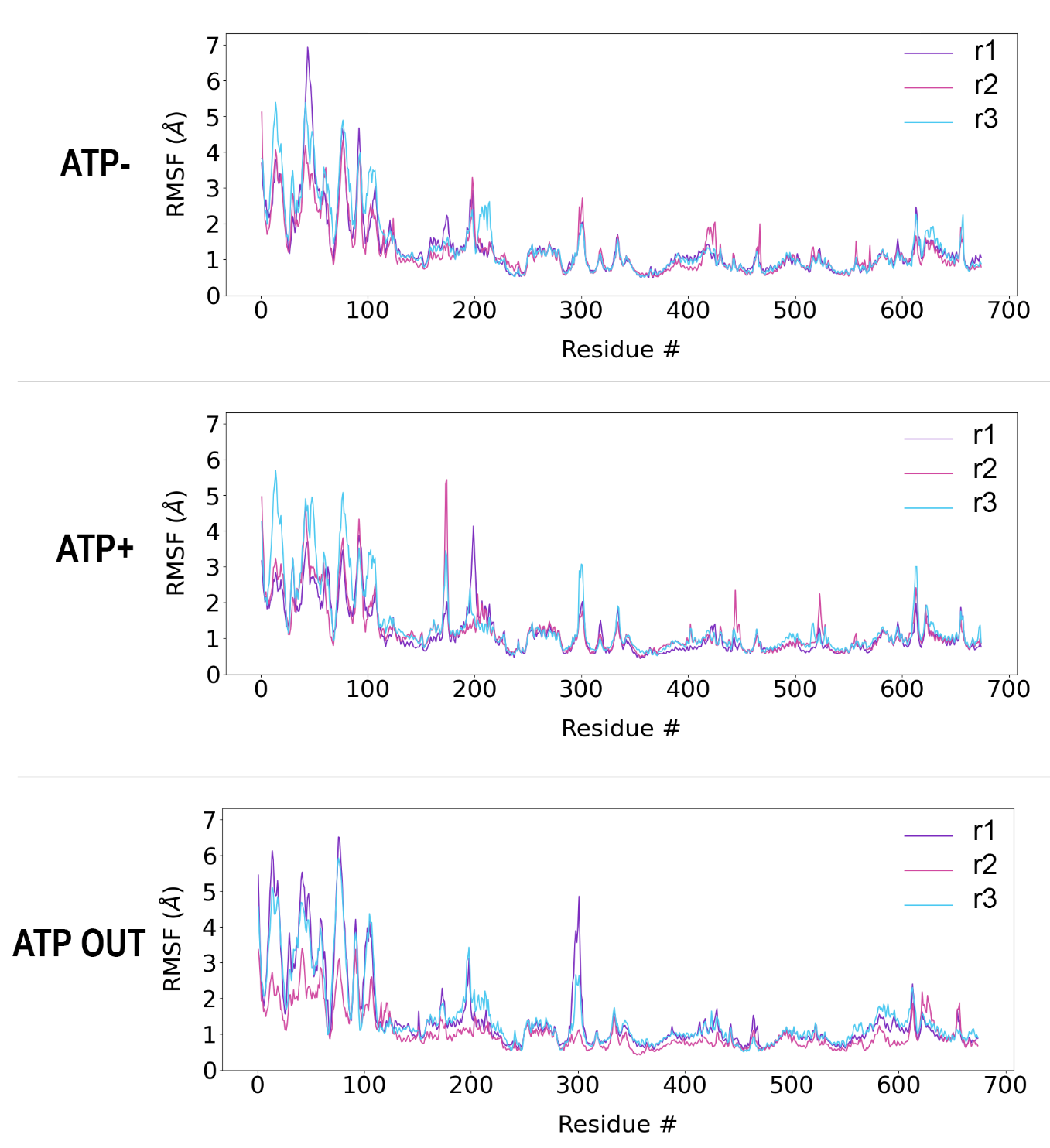

**Root-Mean-Square-Fluctuations (RMSF) per residue in CHARMM36m-Li-Merz simulations^2^.**

**Supplementary Table 3.**

| **Won *et al.* CHARMM ion parameterization** | | | | | | | |
| --- | --- | --- | --- | --- | --- | --- | --- |
| ***Stable  5-LOX*** | **Atom** | **Nr of frames** | **Distance between residues and Fe^2+^** | | | | **Notes** |
|  |  |  | **Mean (Å)** | **SD (Å)** | **Min (Å)** | **Max (Å)** |  |
| **His368** | Nε2 | 602 | 2.47 | 0.7 | 2.02 | 4.93 | *two-state: coordinated ~2.1 Å & excursions ≥4 Å* |
| **His373** | Nε2 | 602 | 2.17 | 0.06 | 2.00 | 2.44 | *tight, classic anchor.* |
| **His551** | Nε2 | 602 | 2.18 | 0.07 | 2.02 | 2.52 | *tight with mild breathing* |
| **Asn555** | Oδ1 | 602 | 2.05 | 0.07 | 1.90 | 2.34 | *stable polar ligand* |
| **Ile674** | O1  O2 | 602 | 2.02  2.22 | 0.25  0.50 | 1.84  1.84 | 3.46  3.52 | ***monodentate carboxylate*** *occasionally switching O atoms* |

**Stable 5-LOX Fe^2+^ coordination distances observed in CHARMM36m force field MD with Fe (II) ion parameters developed by Won *et al*. ^1^ for stable 5 LOX (300 ns, 2X) show extremely tight and stable coordination.**

**Supplementary Figure 10.**

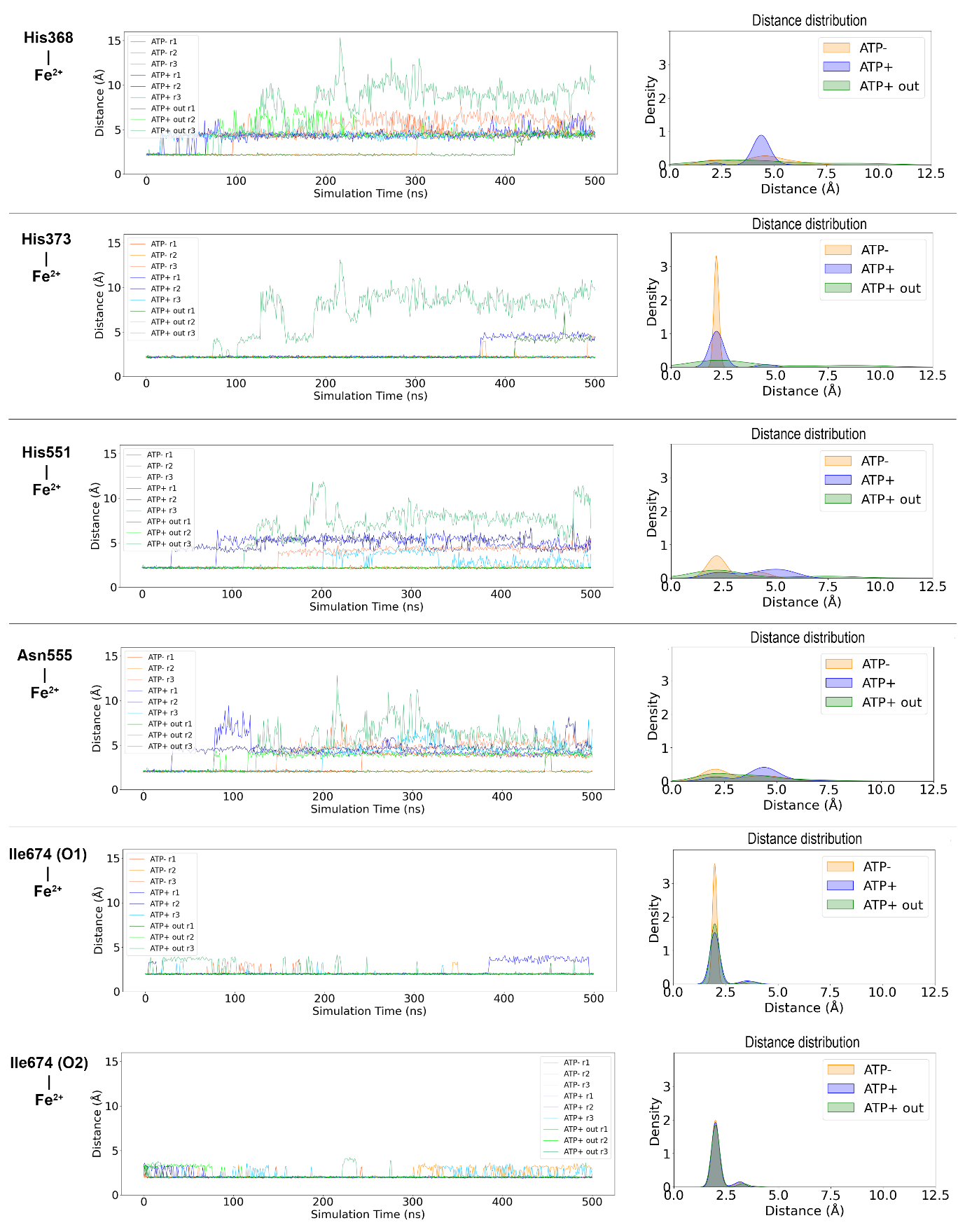

**Time-dependent Fe^2+^–residue distance profiles and corresponding distance distributions for the catalytic iron and its coordinating residues over 500 ns MD simulations using the Won *et al.* ion parameterization.**

**Supplementary Figure 11.**

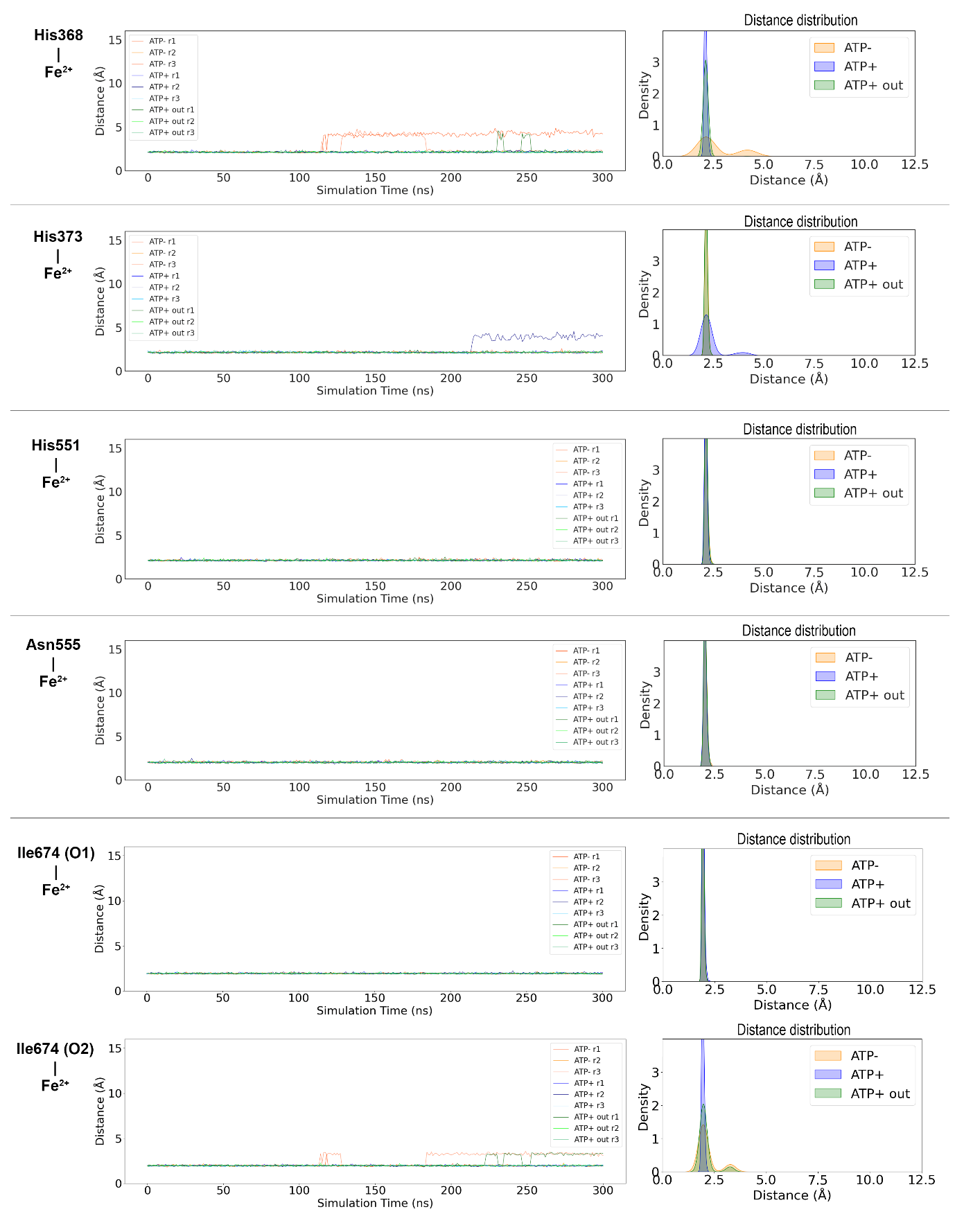

**Time-dependent Fe^2+^–residue distance profiles and corresponding distance distributions for the catalytic iron and its coordinating residues over 500 ns MD simulations using the Li-Merz iron parameterization.**

**Supplementary Table 4.**

| **Won *et al.* ion parameterization** | | | | | | |
| --- | --- | --- | --- | --- | --- | --- |
|  |  | **Nr of frames** | **Distance between residues and Fe^2+^** | | | |
|  |  |  | **Mean (Å)** | **SD (Å)** | **Min (Å)** | **Max (Å)** |
| **His368** | ATP- | 903 | 3.533 | 1.433 | 2.017 | **7.577** |
| Nε2 | ATP+ | 903 | 4.160 | 0.667 | 2.072 | **6.836** |
|  | ATP OUT | 903 | 4.271 | 2.658 | 2.023 | **15.336** |
| **His373** | ATP- | 903 | 2.166 | 0.062 | 1.986 | **2.419** |
| Nε2 | ATP+ | 903 | 2.173 | 0.067 | 2.004 | **2.427** |
|  | ATP OUT | 903 | 3.331 | 2.403 | 2.018 | **13.119** |
| **His551** | ATP- | 903 | 2.492 | 0.736 | 1.994 | **4.689** |
| Nε2 | ATP+ | 903 | 3.944 | 1.441 | 2.002 | **6.435** |
|  | ATP OUT | 903 | 3.202 | 2.149 | 1.972 | **11.879** |
| **Asn555** | ATP- | 903 | 2.636 | 1.114 | 1.831 | **7.691** |
| Oδ1 | ATP+ | 903 | 3.744 | 1.368 | 1.910 | **9.438** |
|  | ATP OUT | 903 | 3.223 | 1.768 | 1.893 | **12.820** |
| **Ile674** | ATP- | 903 | 2.019 | 0.213 | 1.827 | **3.573** |
| O_1_ | ATP+ | 903 | 2.010 | 0.199 | 1.844 | **3.545** |
|  | ATP OUT | 903 | 2.141 | 0.503 | 1.847 | **4.106** |
| **Ile674** | ATP- | 903 | 2.001 | 0.152 | 1.856 | **3.452** |
| O_2_ | ATP+ | 903 | 2.104 | 0.357 | 1.851 | **3.574** |
|  | ATP OUT | 903 | 2.160 | 0.478 | 1.830 | **4.210** |
| **Li-Merz iron parameterization** | | | | | | |
|  |  | **Nr of frames** | **Distance between residues and Fe^2+^** | | | |
|  |  |  | **Mean (Å)** | **SD (Å)** | **Min (Å)** | **Max (Å)** |
| **His368** | ATP- | 903 | 2.679 | 0.918 | 1.951 | **4.872** |
| Nε2 | ATP+ | 903 | 2.118 | 0.065 | 1.960 | **2.448** |
|  | ATP OUT | 903 | 2.139 | 0.220 | 1.958 | **4.568** |
| **His373** | ATP- | 903 | 2.144 | 0.069 | 1.986 | **2.567** |
| Nε2 | ATP+ | 903 | 2.315 | 0.535 | 1.955 | **4.582** |
|  | ATP OUT | 903 | 2.143 | 0.067 | 1.973 | **2.430** |
| **His551** | ATP- | 903 | 2.140 | 0.074 | 1.970 | **2.524** |
| Nε2 | ATP+ | 903 | 2.128 | 0.063 | 1.957 | **2.479** |
|  | ATP OUT | 903 | 2.142 | 0.070 | 1.985 | **2.531** |
| **Asn555** | ATP- | 903 | 2.043 | 0.075 | 1.871 | **2.354** |
| Oδ1 | ATP+ | 903 | 2.029 | 0.074 | 1.833 | **2.510** |
|  | ATP OUT | 903 | 2.038 | 0.073 | 1.860 | **2.433** |
| **Ile674** | ATP- | 903 | 1.932 | 0.050 | 1.790 | **2.157** |
| O1 | ATP+ | 903 | 1.944 | 0.056 | 1.822 | **2.243** |
|  | ATP OUT | 903 | 1.927 | 0.049 | 1.810 | **2.119** |
| **Ile674** | ATP- | 903 | 2.155 | 0.465 | 1.799 | **3.559** |
| O2 | ATP+ | 903 | 1.958 | 0.057 | 1.803 | **2.164** |
|  | ATP OUT | 903 | 2.068 | 0.343 | 1.834 | **3.448** |
| *See related Supplementary Figure 10 and 11.* | | | | | | |

**Comparison of Fe(II) coordination distances over 900ns (300 ns, 3X) for simulations with the Won^1^ versus Li-Merz^2^ parameters for iron.** The Won FF keeps the iron ≈ 2-2.5 for WT simulations in absence of ATP, for all residues but the replaceable His368, while in presence of ATP displays shifts to the second coordination shell for several residues. Li-Merz parameters resemble CHARMM36m-Won simulations of Stable 5-LOX with clear breathing or ATP-induced changes at the catalytic site (Supplementary Table 3).

**Supplementary Table 5.**

| **Won *et al*. iron parameterization** | | | | | | |
| --- | --- | --- | --- | --- | --- | --- |
|  |  | **Nr of**  **frames** | **Distance between residues and Fe^2+^** | | | |
|  |  |  | **Mean (Å)** | **SD (Å)** | **Min (Å)** | **Max (Å)** |
| **His368** | ATP- | 1503 | 4.150 | 1.457 | 2.017 | 7.577 |
| Nε2 | ATP+ | 1503 | 4.310 | 0.621 | 2.072 | 6.836 |
|  | ATP OUT | 1503 | 4.773 | 2.741 | 2.023 | 15.336 |
| **His373** | ATP- | 1503 | 2.183 | 0.203 | 1.986 | 4.704 |
| Nε2 | ATP+ | 1503 | 2.374 | 0.666 | 2.004 | 5.107 |
|  | ATP OUT | 1503 | 3.855 | 2.728 | 2.018 | 13.119 |
| **His551** | ATP- | 1503 | 2.659 | 0.894 | 1.994 | 5.797 |
| Nε2 | ATP+ | 1503 | 4.082 | 1.344 | 2.002 | 7.708 |
|  | ATP OUT | 1503 | 3.555 | 2.457 | 1.972 | 11.879 |
| **Asn555** | ATP- | 1503 | 3.061 | 1.299 | 1.831 | 7.694 |
| Oδ1 | ATP+ | 1503 | 4.068 | 1.218 | 1.910 | 9.438 |
|  | ATP OUT | 1503 | 3.527 | 1.786 | 1.862 | 12.820 |
| **Ile674** | ATP- | 1503 | 2.006 | 0.187 | 1.822 | 3.573 |
| O1 | ATP+ | 1503 | 2.121 | 0.461 | 1.842 | 4.090 |
|  | ATP OUT | 1503 | 2.081 | 0.403 | 1.847 | 4.106 |
| **Ile674** | ATP- | 1503 | 2.093 | 0.346 | 1.856 | 3.602 |
| O2 | ATP+ | 1503 | 2.101 | 0.351 | 1.850 | 3.574 |
|  | ATP OUT | 1503 | 2.092 | 0.383 | 1.830 | 4.210 |

**Extended molecular dynamics simulations (500 ns, 3X) with CHARMM-Won parameterization^1^.**

**Supplementary Figure 12.**

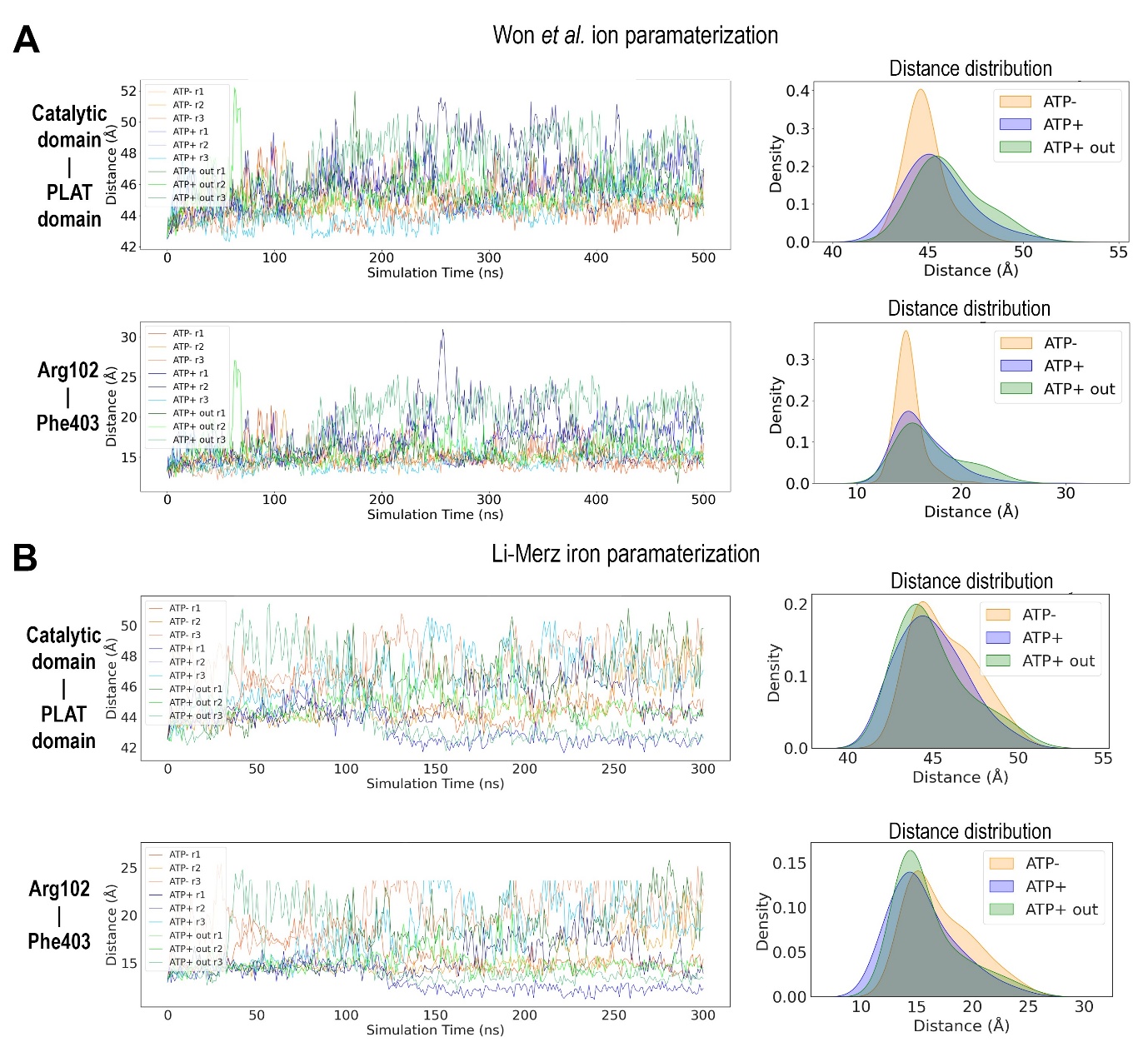

**Time-resolved distances (left) and corresponding distance distributions (right) for (i) the catalytic–PLAT domain separation and (ii) the Arg102–Phe403 pair across ATP–, ATP+, and ATP+out simulations using Won et al.^1^ (A) and Li-Merz^2^ (B) ion parameterization.** The comparison highlights Fe^2+^-parameter-dependent differences in distance fluctuations and sampled conformational states.

**Supplementary Table 6.**

| **Won *et al.* ion parameterization** | | | | | | |
| --- | --- | --- | --- | --- | --- | --- |
|  |  | **Nr of frames (n)** | **Distance between domains or residues** | | | |
|  |  |  | **Mean (Å)** | **SD (Å)** | **Min (Å)** | **Max (Å)** |
| **Catalytic vs**  **PLAT domain** | ATP- | 1503 | 44.835 | 1.038 | 42.580 | 48.794 |
|  | ATP+ | 1503 | 45.533 | 1.710 | 42.317 | 51.583 |
|  | ATP OUT | 1503 | 46.141 | 1.707 | 42.563 | 52.214 |
| **Arg102**  **Vs**  **Phe403** | ATP- | 1503 | 14.941 | 1.223 | 12.120 | 21.457 |
|  | ATP+ | 1503 | 16.220 | 2.489 | 12.317 | 30.978 |
|  | ATP OUT | 1503 | 17.011 | 2.968 | 11.685 | 27.118 |
| **Li-Merz iron parameterization** | | | | | | |
|  |  | **Nr of frames (n)** | **Distance between domains or residues** | | | |
|  |  |  | **Mean (Å)** | **SD (Å)** | **Min (Å)** | **Max (Å)** |
| **Catalytic vs**  **PLAT domain** | ATP- | 903 | 45.706 | 1.799 | 42.522 | 50.794 |
|  | ATP+ | 903 | 44.965 | 1.922 | 41.638 | 50.586 |
|  | ATP OUT | 903 | 44.965 | 2.069 | 41.786 | 51.435 |
| **Arg102**  **Vs**  **Phe403** | ATP- | 903 | 17.138 | 2.883 | 13.241 | 25.994 |
|  | ATP+ | 903 | 15.629 | 2.951 | 11.162 | 26.884 |
|  | ATP OUT | 903 | 16.051 | 2.982 | 12.125 | 26.355 |
| *See related Supplementary Figure 12.* | | | | | | |

**Distance statistics for the catalytic and PLAT domain and Arg102–Phe403 decoupling under the Won *et al.*^1^ and Li–Merz^2^ ion parameterizations.**

**Supplementary Figure 13.**

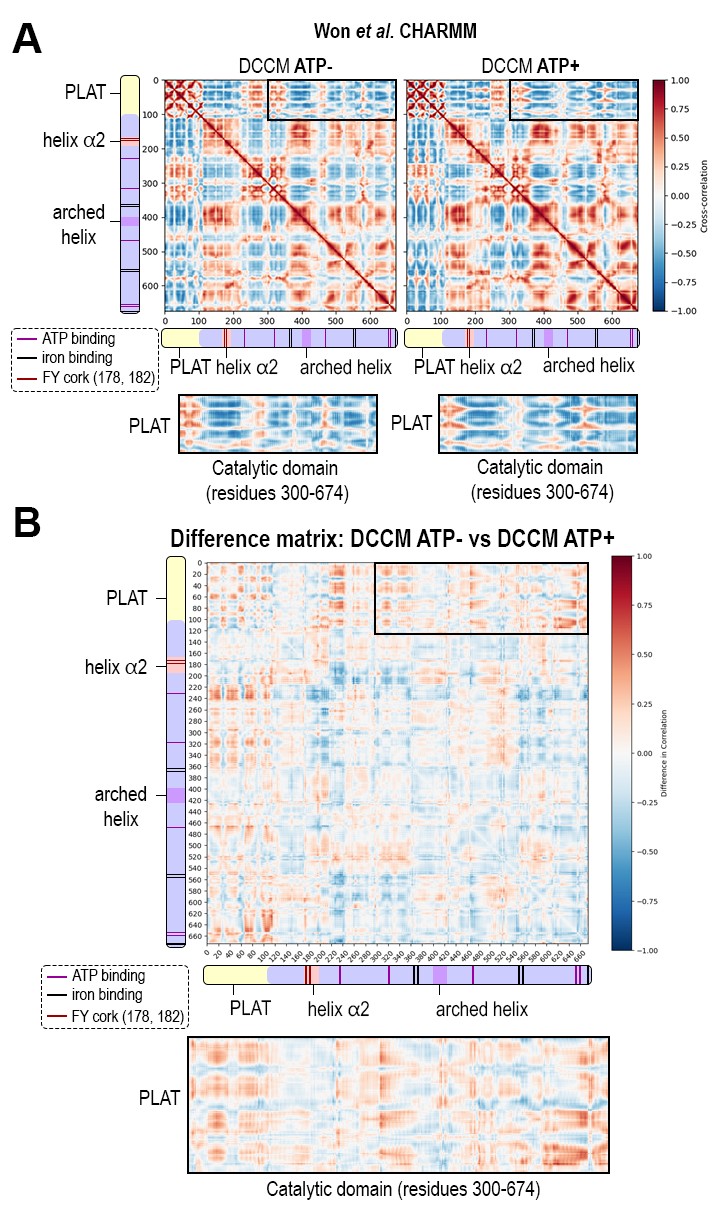

**Dynamic residue–residue cross-correlation matrices for ATP– and ATP+ simulations using the Won ion parameterization. A** Cross-correlation matrices (DCCMs) for ATP– and ATP+ trajectories and **B** the difference matrix (ATP+ – ATP–) highlighting regions where correlations decrease (blue) or increase (red) upon ATP binding. The pattern reveals a net loss of correlated motions between the PLAT and catalytic domains, consistent with enhanced interdomain decoupling and increased maximal opening values (see Supplementary Figure 13).

**Supplementary Figure 14.**

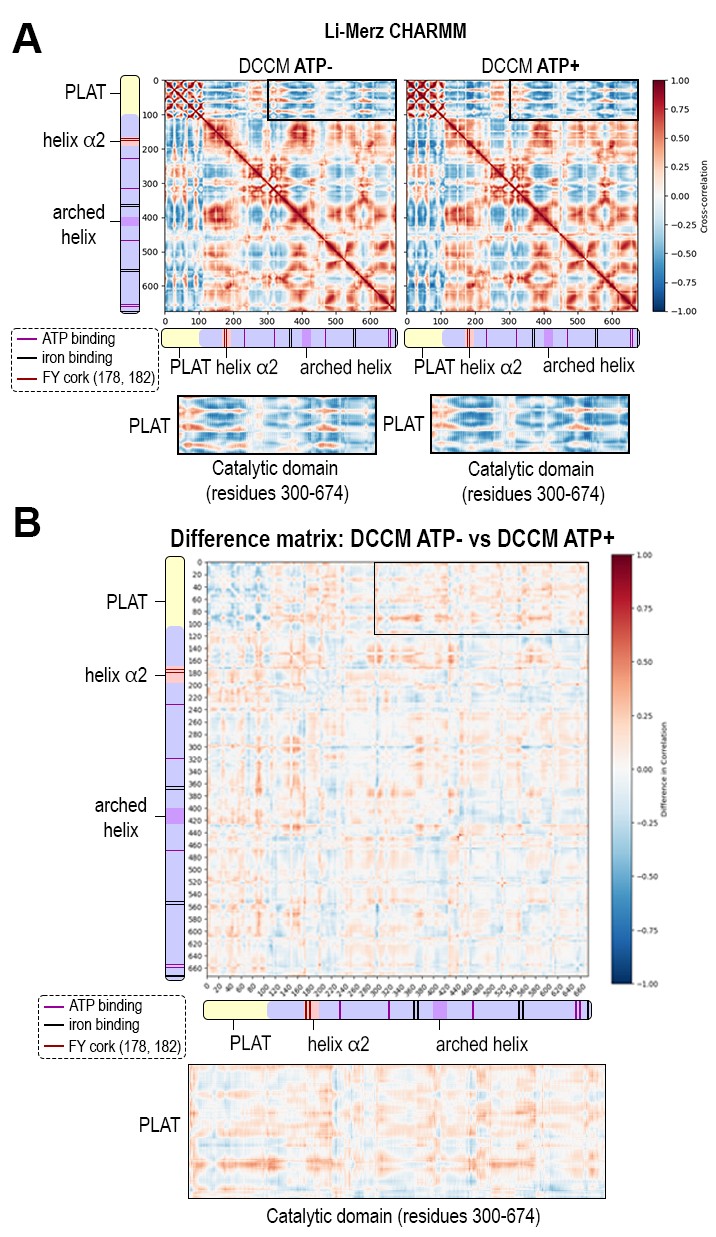

**Dynamic residue–residue cross-correlation matrices for ATP– and ATP+ simulations using the Li-Merz iron parameterization. A** Cross-correlation matrices (DCCMs) for ATP– and ATP+ trajectories and **B** the difference matrix (ATP+ – ATP–) highlighting correlation changes upon ATP binding. Blue regions (negative correlations) and diminished red regions (positive correlations) still indicate interdomain decoupling, but the magnitude of ATP-induced changes is smaller and less coherent across the domain interface compared to the Won approach (see Supplementary Figure 13).

**Supplementary Table 7.**

| **Canonical 5-LOX Fe^2+^ coordination with expected average distances** | | | | |
| --- | --- | --- | --- | --- |
| **Ligand**  **(residue)** | **Atom (donor)** | **Expected** | **Typical range** | **Notes** |
| **His368** | Nε2 | ~2.15 | 2.05–2.40 | Labile / non-essential ligand; may disengage without loss of metal. |
| **His373** | Nε2 | ~2.15 | 2.05–2.25 | Stable axial histidine anchor. |
| **His551** | Nε2 | ~2.05 | 2.05–2.25 | Stable equatorial histidine anchor. |
| **Asn555** | Oδ1 | ~2.05 | 2.00–2.20 | Polar tuning ligand; generally coordinated but more flexible than His. |
| **Ile674** | Terminal COO^-^ O_1_  Terminal COO^-^ O_2_ | ~2.70 | 1.95–2.20  2.5-3.5 | Primary C-terminal carboxylate ligand; usually the main coordinating O.  Typically, non-coordinating; can transiently swap with O₁. |
| **H₂O / OH⁻ (6th site)** | O(wat) | ~2.20 | 2.10–2.40 | Occupancy/state-dependent; often exchanged |

**Canonical 5-LOX Fe^2+^ coordination with expected/theoretical average distances derived from Bazayeva *et al.* 2024^3^.**

**Supplementary Figure 15.**

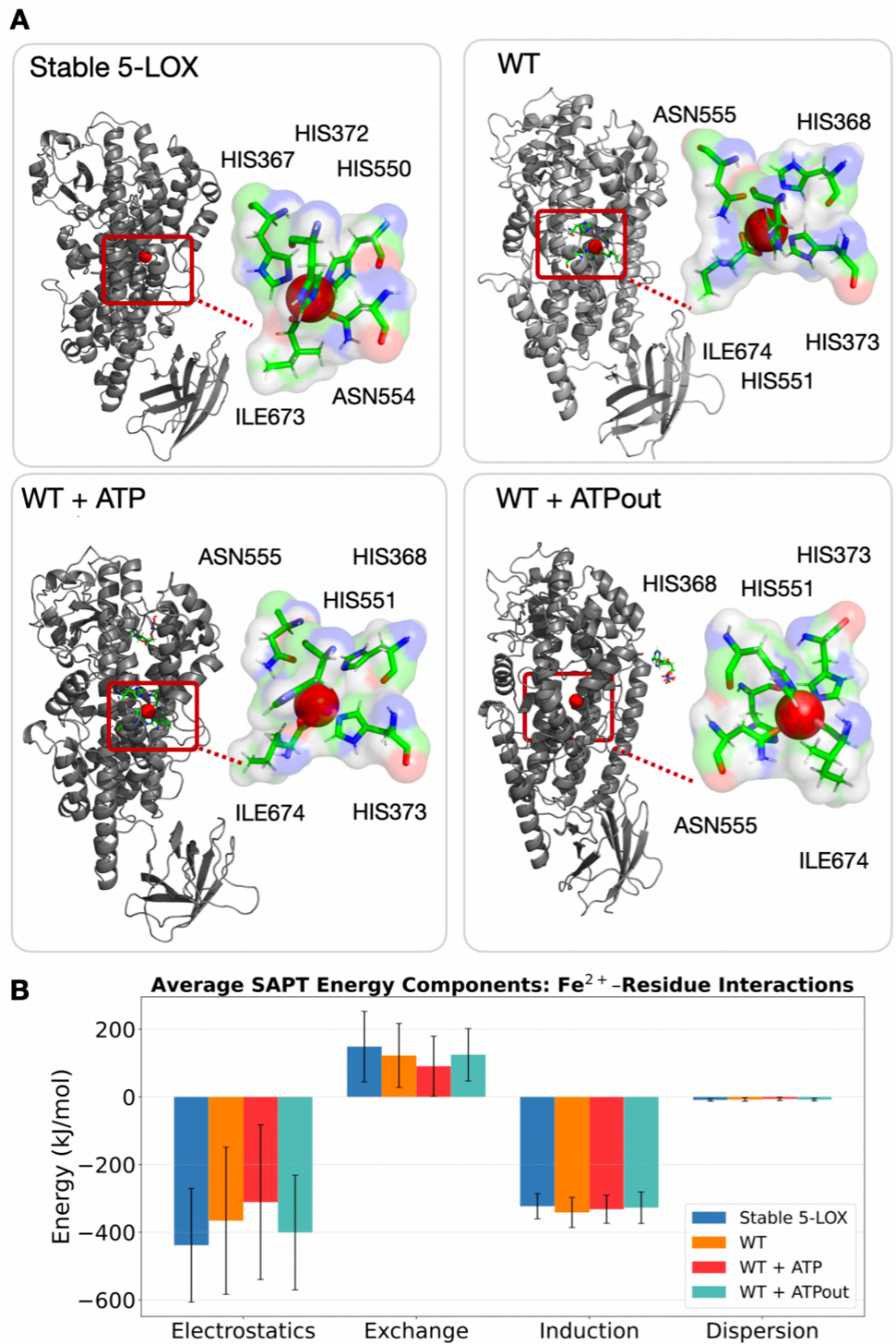

**Quantum-mechanical energy analysis of Fe^2+^ coordination in 5-LOX. A** Representative Fe²⁺-coordination geometries within the catalytic site, highlighting interactions with His368, Asn555, His373, His551, and Ile674 (stable variant: His367, His372, His550, Asn554, and Ile673). The structures correspond to one of the five representative conformations obtained from molecular dynamics simulations for the following systems: the stable variant, the wild-type enzyme, the wild-type enzyme with ATP bound, and the wild-type enzyme in the outward-facing ATP-bound state. **B** Induction, electrostatic, exchange, and dispersion components of Fe²⁺–residue interactions averaged over five representative conformations (in kJ mol⁻¹), shown as mean ± SD over five representative conformations for each system. Interactions were computed between Fe²⁺ and the coordinating donor atoms (His368, His373, His551, Asn555, and Ile674; stable variant: His367, His372, His550, Asn554, and Ile673). Electrostatic and induction terms dominate the overall stabilization across all systems.

**Supplementary Table 8.**

| **Residue** | **Atom** | **Distance (Å)** | **Distance (Å)** | **Distance (Å)** | **Distance (Å)** |
| --- | --- | --- | --- | --- | --- |
|  |  | **WT** | **WT + ATP** | **WT + ATP** | **Stable 5-LOX** |
| **His368** | Nε2 | 3.48 ± 1.22 | 3.46 ± 1.21 | 2.93 ± 1.08 | 2.25 ± 0.12 |
| **His373** | Nε2 | 2.14 ± 0.04 | 2.19 ± 0.09 | 2.21 ± 0.07 | 2.12 ± 0.04 |
| **His551** | Nε2 | 2.11 ± 0.07 | 3.64 ± 1.36 | 2.15 ± 0.07 | 2.19 ± 0.09 |
| **Asn555** | Oδ1 | 2.97 ± 0.88 | 3.29 ± 1.09 | 2.07 ± 0.08 | 2.03 ± 0.03 |
| **Ile674** | O | 2.03 ± 0.05 | 2.00 ± 0.06 | 2.13 ± 0.40 | 2.18 ± 0.46 |

**Detailed Fe²⁺–amino acid distances (Å) obtained from representative low-coordination snapshots from CHARMM36m–Won simulations used for QM SAPT energy decomposition analyses.** Distances correspond to the mean separation between the indicated residue atom and the iron center and are reported as mean ± SD over five conformations (see Supplementary Fig. 15).

**Supplementary Figure 16.**

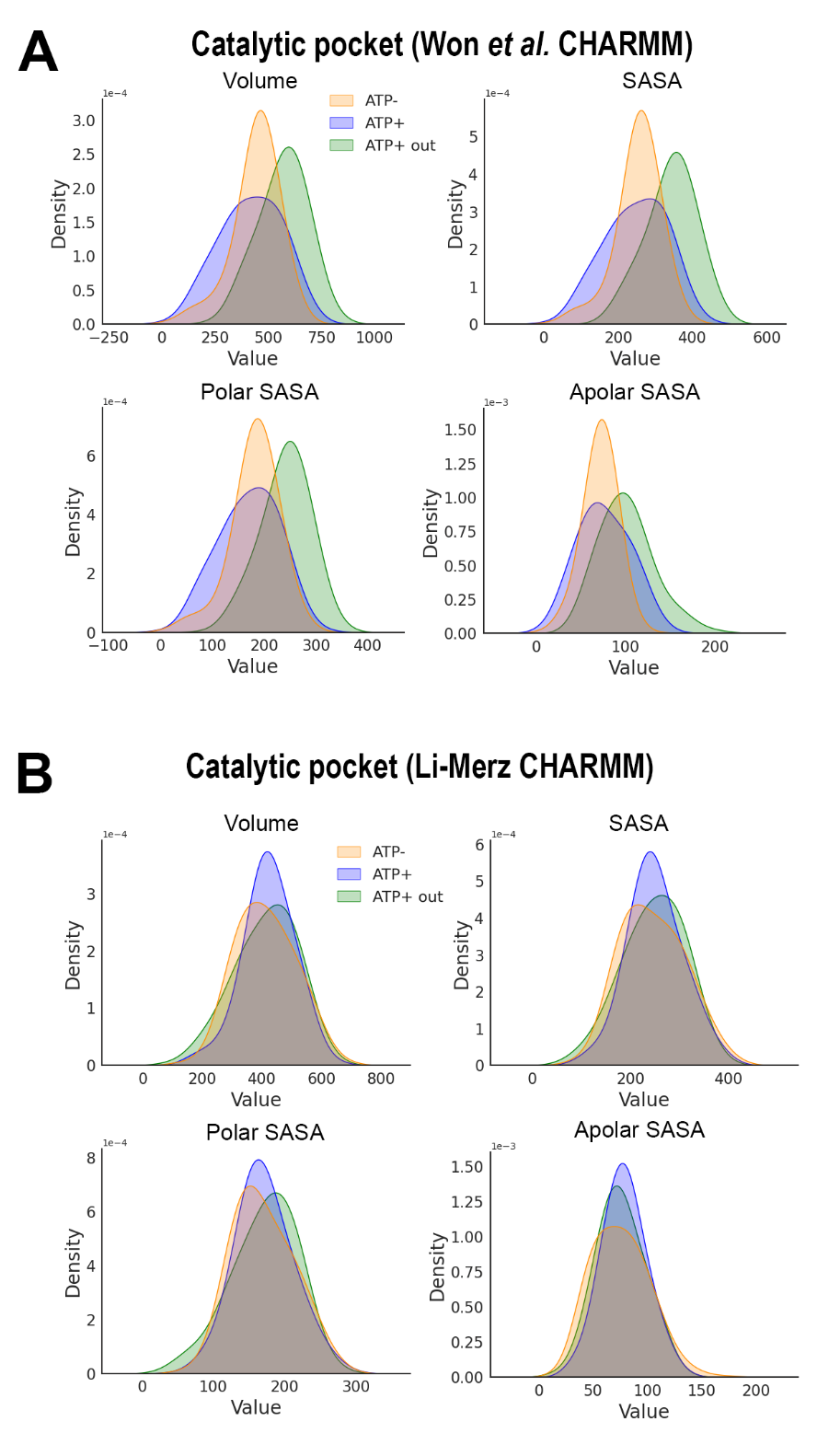

**Main features of the catalytic pocket in CHARMM-Won versus Li-Merz parameters.**

**SUPPLEMENTARY DISCUSSION**

**SAPT Energy Decomposition of Fe²⁺–Ligand Interactions in Low coordination states**

Symmetry-adapted perturbation theory (SAPT0/def2-SVP) calculations were performed on representative low-coordination MD snapshots from CHARMM36m-Won MD (see Methods and Supp. Table 8) for the WT 5-LOX active site under ATP– and ATP+ conditions, in order to evaluate the physical energy components stabilizing Fe²⁺–ligand interactions. The decomposition into electrostatic, induction (polarization + charge transfer), exchange (Pauli repulsion), and dispersion terms provides a quantitative description of metal–ligand stabilization and its modulation by ATP binding (Supp. Fig. 15) From the most probable conformations sampled in the MD simulations, the SAPT energy decomposition reveals the following behaviors for the Fe²⁺ ligands.

For **His551**, the Fe–NE2 interaction in the ATP– frame was dominated by electrostatic (−388.3 kJ mol⁻¹) and induction (−328.1 kJ mol⁻¹) contributions, opposed by exchange repulsion (+128.4 kJ mol⁻¹). Upon ATP binding, this residue became more labile, with the electrostatic term decreasing sharply to −28.3 kJ mol⁻¹ while the induction component remained strongly attractive (−281.4 kJ mol⁻¹), consistent with partial detachment yet sustained polarization coupling to Fe²⁺.

For **His373**, a core anchoring ligand, the electrostatic (−326.2 kJ mol⁻¹) and induction (−333.2 kJ mol⁻¹) energies in the ATP– structure confirm a robust interaction with the metal center. In the ATP+ frame, these terms remain comparably large (−310.0 and −317.2 kJ mol⁻¹, respectively), indicating a persistent and strongly stabilizing coordination even under the more flexible ATP-bound configuration.

The terminal **Ile674** carboxylate exhibits the largest stabilizing energies in both states, with electrostatics of −580.1 kJ mol⁻¹ and induction of −424.9 kJ mol⁻¹ in the ATP– frame, further strengthened to −687.5 and −420.8 kJ mol⁻¹, respectively, in the ATP+ frame. This residue thus remains a key anchor for Fe²⁺ across all conformations.

By contrast, **His368** and **Asn555** show weaker electrostatics (−40 to −80 kJ mol⁻¹) but substantial induction energies (−315 to −365 kJ mol⁻¹), consistent with labile or second-shell coordination behavior. Their longer Fe–donor distances (3.5–4.5 Å) place them in the weak-coordination regime described for Fe sites^3^, where polarization and charge-transfer interactions dominate over direct bonding. Notably, this agrees with experimental data indicating their replaceable nature^4, 5, 6, 7^.

Overall, the SAPT analysis confirms that Fe²⁺ stabilization in 5-LOX arises primarily from **induction and electrostatic components**, with His373 and Ile674 providing persistent first-shell anchoring and His368, His551, and Asn555 contributing flexible, second-shell polarization interactions. The total interaction energies remain attractive under both ATP– and ATP+ conditions, supporting the interpretation that the ATP-induced low-coordination geometries observed in MD correspond to **energetically plausible, partially relaxed and transient states** that could play a role in the accommodation of the substrate in the binding site.

**MATERIALS AND METHODS**

**Supplementary Table 9.**

| **Chemical / Material** | **Abbreviation** | **Company** |
| --- | --- | --- |
| 13*S*-hydroperoxy-octadecaenoic acid | 13*S*-HpODE | Cayman Chemical |
| 17*S*-hydroxy-docosatetraenoic acid | 17*S*-HDoTE | Cayman Chemical |
| 5*S*-hydroxy-eicosatetraenoic acid | 5*S*-HETE | Cayman Chemical |
| 5*S*-hydroperoxy-eicosatetraenoic acid | 5*S*-HpETE | Cayman Chemical |
| Acetonitrile | ACN | Merck |
| Arachidonic acid | AA | Cayman Chemical |
| Adenosine 5’-monophosphate | AMP | Merck |
| Ampicillin | - | Merck |
| Adenosine 5’-triphosphate | ATP | Merck |
| Benzonase® Nuclease | - | Merck |
| *E. coli* BL21(DE3) competent cells for expression | BL21(DE3) | Thermo Fisher Scientific |
| cOmplete™ EDTA-free protease inhibitor cocktail | - | Roche |
| *E. coli* DH5α competent cells for cloning | DH5α | Thermo Fisher Scientific |
| *E. coli* DH10β competent cells for cloning | DH10β | Thermo Fisher Scientific |
| Formic acid | FA | Merck |
| Hen egg lysozyme | Lysozyme | Roche |
| Hydrogen chloride | HCl | Merck |
| Isopropyl β-D-1-thiogalactopyranoside | IPTG | Merck |
| Leukotriene A_4_ | LTA_4_ | Med Chem 101 |
| Methanol | MeOH | Merck |
| Tris(hydroxymethyl)aminomethane (base and acetate) | Tris | Merck |
| Sodium chloride | NaCl | Merck |
| StrepTactin XT 4Flow resin |  | IBA Lifesciences |
| Terrific Broth medium (tryptone, yeast extract, glycerol, mono- and dibasic potassium phosphate) | TB medium | Merck |

**Materials used in the experimental part.**

**Supplementary Table 10.**

| **Mutation** |  | **Primer sequence (5’-3’)** |
| --- | --- | --- |
| K320L | F | GCGAACctAATCGTGCCGATC |
|  | R | CACGATTagGTTCGCCAGGTTC |
| Y468A | F | ACATCCCGgcCTATTTTTATCGTG |
|  | R | AAATAGgcCGGGATGTCTTCTTTG |
| K656L | F | AAGAAActGCAACTGCCGTATTATTATC |
|  | R | CAGTTGCagTTTCTTGTTACGTTCC |
| Q657L | F | GAAAAAGCtACTGCCGTATTATTATCTGAG |
|  | R | GGCAGTaGCTTTTTCTTGTTACGTTC |
| F – a forward primer, R – a reverse primer. | | |

**Primers used for the whole plasmid PCR to introduce amino acid replacements in the ATP binding pocket of 5-LOX.** Mutated codons are shown in lowercase.

**Supplementary Table 11.**

| **MD minimization parameters** | |
| --- | --- |
| Constraints | h-bonds (LINCS algorithm) |
| Cutoff scheme | Verlet |
| Neighbor list update | every 10 steps |
| van der Waals interactions | Force-switch method with a cutoff of 1.2 nm, switching at 1.0 nm |
| Electrostatics | Particle Mesh Ewald (PME) with a cutoff of 1.2 nm |
| **MD equilibration parameters** | |
| Integrator | Molecular dynamics (md) |
| Timestep (dt) | 0.001 ps |
| Simulation length (nsteps) | 125 000 steps (125 ps) |
| *Output frequencies* | |
| Compressed trajectory output (nstxout-compressed) | every 5000 steps |
| Energy output (nstenergy) | every 1000 steps |
| Log output (nstlog) | every 1000 steps |
| *Cutoff settings* | |
| Cutoff scheme: | Verlet |
| Neighbor list update (nstlist) | every 20 steps |
| van der Waals interactions | Force-switch, cutoff 1.2 nm, switching at 1.0 nm |
| Electrostatics | Particle Mesh Ewald (PME), cutoff 1.2 nm |
| Thermostat (tcoupl) | v-rescale |
| Temperature groups (tc_grps) | SOLU, SOLV |
| Temperature coupling time constant (tau_t) | 1.0 ps for both groups |
| Reference temperature (ref_t) | 303.15 K for both groups |
| Center of mass motion removal (nstcomm) | every 100 steps |
| Mode (comm_mode): | linear |
| Center of mass groups (comm_grps): | SOLU, SOLV |
| Velocity generation (gen_vel): | yes |
| Generation temperature (gen_temp): | 303.15 K |
| Random seed (gen_seed): | -1 (randomized) |
| *Position restraints* | |
| Backbone | POSRES_FC_BB=400.0 |
| Side chains | POSRES_FC_SC=40.0 |
| **MD production parameters** | |
| Integrator | Molecular dynamics (md) |
| Timestep (dt) | 0.002 ps |
| *Output frequencies* | |
| Compressed trajectory output (nstxout-compressed): | every 50,000 steps |
| Energy calculations (nstcalcenergy) | every 100 steps |
| Energy output (nstenergy): | every 1,000 steps |
| Log output (nstlog): | every 1,000 steps |
| *Cutoff settings* | |
| Cutoff scheme | Verlet |
| Neighbor list update (nstlist) | every 20 steps |
| van der Waals interactions | Force-switch, cutoff 1.2 nm, switching at 1.0 nm |
| Electrostatics | Particle Mesh Ewald (PME), cutoff 1.2 nm |
| Thermostat (tcoupl) | v-rescale |
| Temperature groups (tc_grps) | SOLU, SOLV |
| Temperature coupling time constant (tau_t) | 1.0 ps for both groups |
| Reference temperature (ref_t) | 303.15 K for both groups |
| Barostat (pcoupl) | C-rescale |
| Pressure coupling type (pcoupltype) | isotropic |
| Pressure coupling time constant (tau_p) | 5.0 ps |
| Compressibility (compressibility) | 4.5e-5 |
| Reference pressure (ref_p) | 1.0 bar |
| *Constraints* | |
| h-bonds | LINCS algorithm |
| Constraint algorithm | LINCS |
| Center of mass motion removal (nstcomm) | every 100 steps |
| Mode (comm_mode) | linear |
| Center of mass groups (comm_grps) | SOLU, SOLV |
| Velocity generation (gen_vel) | yes |
| Generation temperature (gen_temp) | 303.15 K |
| Random seed (gen_seed) | -1 (randomized) |
| *All the parameters were used for both the Won et al.^1^ and Li-Merz^2^ approaches.* | |

**Supplementary Methods 1**

**Quantum-mechanical energy decomposition**

To evaluate the energetic plausibility of low coordination Fe²⁺–ligand configurations observed in Won simulations in presence of ATP, representative structures were subjected to symmetry-adapted perturbation theory (SAPT0) calculations using the def2-SVP basis set. Frames were selected from the most probable low coordination conformational clusters —where one or more Fe–donor distances extended to 3.5-5.0 Å—identified with MDAnalysis, focusing on snapshots in which all Fe–ligand distances (His368, His373, His551, Asn555, and Ile674) remained within ±0.05 Å of their mean values for each condition. SAPT0 calculations were performed using Psi4 (v1.10)^8, 9^, decomposing the total Fe²⁺–amino acid interaction energy into electrostatic, induction (polarization and charge transfer), exchange (Pauli repulsion), and dispersion components. The goal of this analysis was to quantify how the magnitude and character of Fe–ligand interactions change upon ATP binding and to verify that the elongated coordination states remain chemically meaningful. All Fe–ligand interactions exhibited overall attractive (negative) total interaction energies, confirming that even the most relaxed, low-coordination geometries sampled during MD correspond to physically stable and chemically plausible configurations rather than force-field artifacts.

**LIST OF REFERENCES**
